# Raman microscopy-based quantification of the physical properties of intracellular lipids

**DOI:** 10.1101/2020.11.19.389775

**Authors:** Masaaki Uematsu, Takao Shimizu

**Affiliations:** Department of Lipid Signaling, National Center for Global Health and Medicine, 1-21-1 Toyama, Shinjuku-ku, Tokyo 162-8655, Japan; Department of Lipidomics, Graduate School of Medicine, The University of Tokyo, 7-3-1 Hongo, Bunkyo-ku, Tokyo 1130-0033, Japan; Institute of Microbial Chemistry, 3-14-23 Kamiosaki, Shinagawa-ku, Tokyo 141-0021, Japan

**Author notes:** Correspondence should be addressed to Masaaki Uematsu.

## Abstract

The physical properties of lipids, such as viscosity, are homeostatically maintained in cells and are intimately involved in physiological roles. Measurement of the physical properties of plasma membranes has been achieved primarily through chemical or genetically encoded fluorescent probes. However, as most probes target plasma membranes, physical properties of lipids in intracellular organelles, including lipid droplets (LDs) are yet to be analyzed. Here, we present a novel Raman microscopy-based approach for quantifying the physical properties of intracellular lipids under deuterium-labeled fatty acid treatment conditions. Focusing on the fact that Raman spectra of carbon-deuterium vibration are altered depending on the surrounding lipid species, we quantitatively represented the physical properties of lipids as the gauche/trans conformational ratio of the introduced labeled fatty acids, which can be used as an indicator of viscosity. Intracellular Raman imaging revealed that the gauche/trans ratio of cytosolic regions was robustly preserved against perturbations attempting to alter the lipid composition. This was likely due to LDs functioning as a buffer against excess gauche/trans ratio. Our novel approach enables the observation of the physical properties of organelle lipids, which is difficult to perform with conventional probes, and is useful for quantitative assessment of the subcellular lipid environment.

## INTRODUCTION

Forming the boundaries of cells and organelles, lipids are a fundamental component of all living organisms, with their physical properties homeostatically maintained inside cells mainly via the tuning of phospholipid acyl chain composition^1,2^. In general, a higher degree of acyl chain unsaturation increases membrane flexibility. The physical properties of lipids are also involved in various physiological roles, such as enzymatic activities and signal induction, through the regulation of membrane protein diffusion or membrane rigidity^3–5^. As such, various probes and methods have been developed to quantify the physical properties of biological membranes at the subcellular level, predominantly through fluorescence techniques^6,7^. However, these methods require the introduction of chemical or genetically encoded fluorescent probes, thereby cannot avoid the artificial effect of introduced probes themselves on the physical properties of the membranes being analyzed. In addition, majority of probes mainly target plasma membranes, and the other intracellular structures are yet to be analyzed. In particular, physical properties of lipid droplets (LDs), which are mainly composed of triglyceride, were hardly quantified nor studied due to the lack of appropriate probes, despite the fact that nearly all eukaryotic cells can create LDs and that their lipid compositions vary in response to external stimuli^8^. Thus, it remains unclear how LDs actually contribute to the physical properties of the intracellular lipid environment, and they are thought to only store excess fatty acids in the form of triglycerides as an energy source^8^.

Raman microscopy is a promising approach to circumvent these problems^9^. As Raman spectra reflect the vibrational modes of molecules, they provide information regarding a molecule’s chemical structure, but they also have a potential to reveal multiple characteristics that may affect a molecule’s vibrational mode. Previous *in vitro* studies have demonstrated that Raman spectra of carbon-hydrogen (C–H) stretch in fatty acids or phospholipids show different shapes depending on viscosity changes caused by altered temperature or pressure^10,11^. However, it has been unknown whether the spectrum of C–H stretch spectrum is sensitive enough to differentiate between the physical properties of lipids under physiological *in vivo* conditions. In addition, as majority of biomolecules contain C–H bonds in their substructures, it is difficult to obtain a pure C–H stretch spectrum for intracellular lipids using Raman microscopy. Furthermore, analytical methods must be developed to convert the observed spectrum into a quantitative value that reflects the physical properties of lipids. A recent report using stimulated Raman scattering (SRS) microscopy demonstrated that C16:0(d31) (fatty acids with sixteen carbons and zero double bonds labeled with thirty-one deuterium atoms) treatment results in the formation of phase-separated endoplasmic reticulum (ER)-associated membrane domains, quantifying the physical properties of lipids inside these domains as the lateral diffusion coefficient^12^. However, calculation of the diffusion coefficient was based on spatial-temporal analysis resulting from C16:0(d31) pulse labeling, thus whether the physical properties of lipids can be quantitatively obtained from spectral transitions is yet to be established.

Here, we present a Raman microscopy-based approach for quantifying the physical properties of subcellular lipids under deuterium-labeled fatty acid treatment using the gauche/trans conformational ratio, which can be used as an indicator of viscosity^13^. This novel approach, that is, using the target lipid itself as a probe, overcame the innate problems of conventional probes that affect the lipid environment, and enabled the investigation of the physical properties of LDs at the same time. Based on experimentally measured reference spectral transitions *in vitro,* and model membrane simulations *in silico*, observed spectra were quantitatively converted into a gauche/trans ratio of labeled fatty acids. Applying this method to intracellular imaging revealed that while the gauche/trans ratio in LDs dynamically changed, it was relatively constant in non-LD regions. This suggested that LDs might function to buffer the intracellular gauche/trans ratio, thereby maintaining the lipid environment in cytoplasmic regions. Our quantitative evaluation will contribute to the understanding of biological functions and regulatory mechanisms associated with the physical state of intracellular lipid environments.

## RESULTS

### Spectra of saturated fatty acids reflect the physical properties of lipids

Raman spectra derived from C–D stretch appear in the silent region (around 1800–2600 cm^−1^), where the spectra from cellular biomolecules are hardly detectable. This means that the spectra of deuterium-labeled fatty acids can be observed with minimal interference from other molecules, even if they are taken up into cells. Therefore, we focused on deuterium-labeled fatty acids, one of the simplest lipids, as candidate probes for measuring the intracellular physical properties of lipids. First, we investigated whether drastic changes in the physical state of lipids, namely, changes in the liquid-solid phase of lipids, are detectable using the Raman spectra of C–D stretch from deuterium-labeled fatty acids. To this end, the standard spectra of deuterium-labeled saturated and unsaturated fatty acids (*in vitro* spectra) were compared with the spectra of each fatty acid from LDs after their incorporation into HeLa cells (*in vivo* spectra) (Fig. 1A and S1A). Saturated and unsaturated fatty acids used in this experiment are in solid (melting temperature > 60°C) and liquid (melting temperature < 14°C) states, respectively, at room temperature, while fatty acids incorporated into cells should be closer to the liquid state, as they are mixed with other endogenous lipids. Therefore, differences between the *in vitro* and *in vivo* spectra of labeled saturated fatty acids were expected to be larger than that of unsaturated fatty acids. Spectra comparisons were qualitatively performed by fitting reconstructed *in vitro* spectra (solid red lines in Fig. 1A and S1A) to *in vivo* spectra (solid black lines in Fig. 1A and S1A); the reconstruction was performed using the weighted sum of *in vitro* spectra, background spectra, and baseline drift (dashed red lines, dashed gray lines, and dashed gray straight lines, respectively, in Fig. 1A and S1A), as previously described^14^. As a result, the spectra of labeled saturated fatty acids (C16:0[d2], C16:0[d31], C18:0[d3], C18:0[d5], and C20:0[d4]) clearly showed poorer fitting than that of labeled unsaturated fatty acids (C18:1[d17], C18:2[d4], C18:3[d14], C18:2[d11], C20:4[d8], C20:4[d11], C20:5[d5], and C22:6[d5]).

**Fig. 1.**
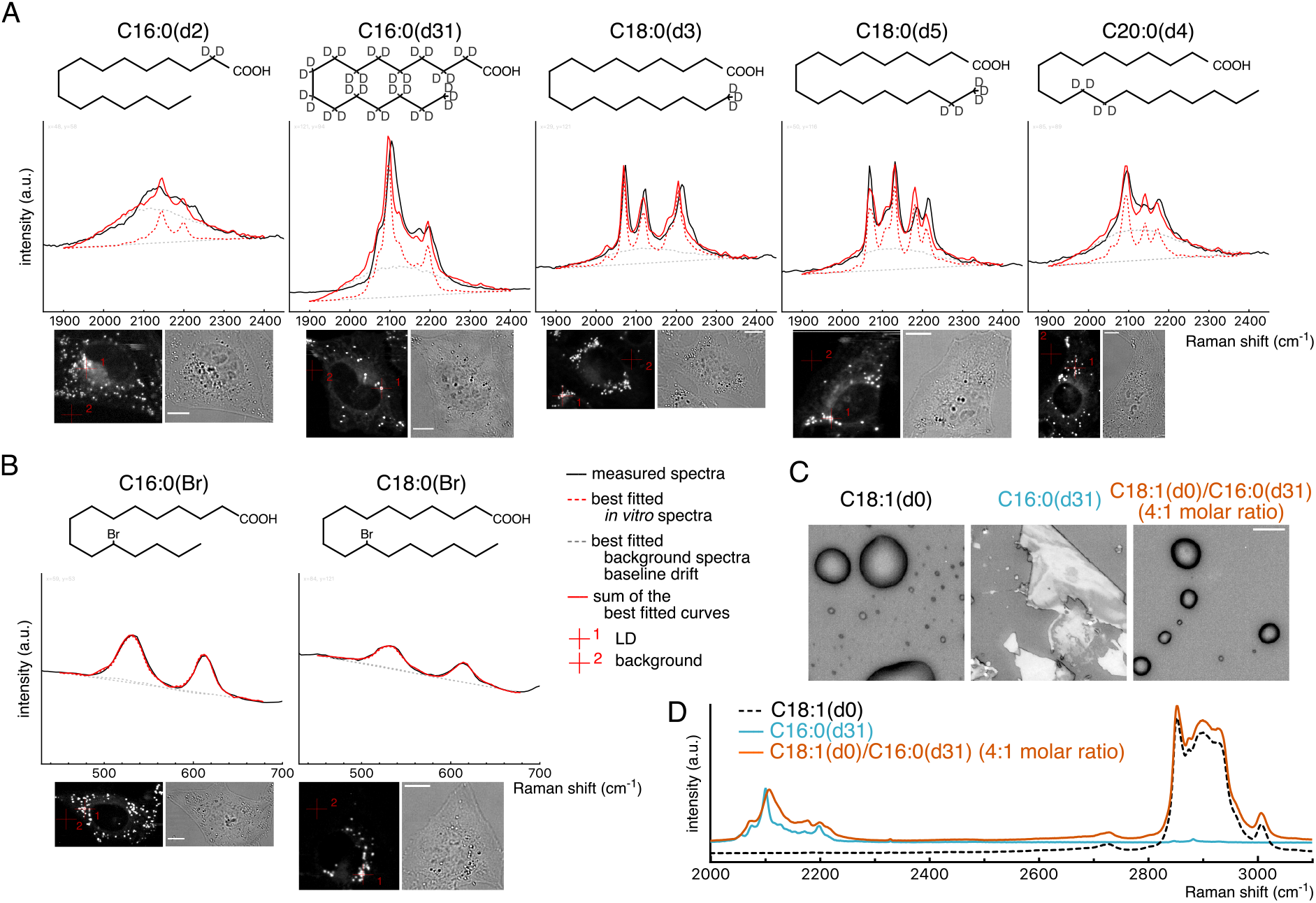
Spectra of saturated fatty acids are affected by the surrounding lipid environment. (**A**) Comparison of Raman spectra of saturated fatty acid between *in vivo* and *in vitro*. HeLa cells treated with indicated deuterium-labeled fatty acids (30 μM) for 24 h were fixed and observed using Raman microscopy. Representative measured spectra at LDs (represented by red crosshair 1 in the bottom images) are shown in black lines, and the results of the best-fitted *in vitro* spectra, background spectra, baseline drifts, and their summations are shown in dashed red lines, dashed gray lines, dashed gray straight lines, and solid red lines, respectively. Spectra at red crosshair 2 in the bottom images were used as the background spectra for the fittings. (**B**) The results of the Raman imaging of Br-labeled fatty acids. The description of the line types of the spectra and the cross hairs in the images are the same as for (A). (**C, D**) Bright field image and Raman spectra of C16:0(d31), C18:1(d0), and their mixture. The Y-axis of each spectra were arbitrarily scaled to make the comparison clearer by equalizing the heights of spectra derived from C–D stretching (C16:0[d31] and the C18:1[d0]/C16:0[d31] mixture) and C–H stretch (C18:1[d0] and the C18:1[d0]/C16:0[d31] mixture). The spectrum of C18:1(d0) (dashed black line) was lowered to avoid overlap. Measurement of multiple cells and samples resulted in the similar results. Scale bars indicate 10 μm.

We also confirmed the correlation between spectral transitions and the solid-liquid state of lipids in another way, using bromine (Br)-labeled saturated fatty acids C16:0(Br) and C18:0(Br), both of which exist in the liquid state at room temperature (Fig. S1B). In contrast to the results of C16:0(d2), C16:0(d31), C18:0(d3), and C18:0(d5), *in vivo* spectra from the carbon-bromine (C–Br) stretch of C16:0(Br) and C18:0(Br), which appears around 533 and 612 cm^−1^, were very similar to *in vitro* spectra (Fig. 1B). These results confirm that *in vivo* and *in vitro* spectral differences represent variances in the physical state of fatty acids. The results also suggest that spectral transitions do not result from the metabolism of incorporated labeled-fatty acids, as some fractions of C16:0(Br) and C18:0(Br) are known to undergo metabolic processes^15^.

Further *in vitro* experiments were performed to determine that the spectral differences represented not only the discontinuous physical state of lipids, that is, solid and liquid states, but also represented continuous physical properties of lipids. For this purpose, we chose C16:0(d31), which displayed the clearest differences between *in vitro* and *in vivo* spectra, with the latter exhibiting a wider maximum peak around 2100 cm^−1^ and a slight right shift of the peak top. While the pure C16:0(d31) *in vitro* were in a solid-like states, a mixture of C18:1(d0) and C16:0(d31) in a 4:1 molar ratio appeared to exist in a liquid-like state, with spectra of the C–D stretch from C16:0(d31) also showing similar transitions to those observed *in vivo* (Fig. 1C and D). Notably, the spectral transitions were not discontinuous, but the gradual changes of the spectra were observed depending on the amount of C18:1(d0) (Fig. S4A). These results indicate that the spectral transitions of C16:0(d31) reflect more than the phase differences, and represent some continuous physical properties of lipids.

### Gauche/trans conformations affect the spectral shape of C16:0(d31)

Next, we investigated what kind of physical properties the spectral transitions represented. Since Raman spectra represent the vibrational modes of molecules, and they can be affected by the conformational state of those molecules. In fatty acids, three interconverting conformational isomers exist for four consecutive carbon atoms (A–B–C–D): one trans and two gauche conformations (Fig. 2A). These isomers correspond to three local energy minima of the four consecutive carbon atoms: when the dihedral angle between planes, defined by two sets of three carbon atoms (A–B–C and B–C–D), is approximately 180 and ± 60 degrees. In the case of phospholipid fatty acyl groups, the ratio of the four consecutive carbon atoms with gauche conformers to those with trans conformers (referred to as the gauche/trans ratio hereafter) increases with temperature^13^. Previous studies also reported that Raman spectra of C–H or C–D stretching from phospholipids, as well as *in vitro* membrane viscosity, can be altered under similar conditions^16^. Therefore, we investigated whether the spectral changes observed in Fig. 1 could be reproduced *in silico* by modulating the gauche/trans conformation. For this, we used density functional theory (DFT), which was developed to perform quantum mechanical modeling to calculate the electronic distribution and vibrational characteristics of compounds. Raman spectra of two types of C16:0(d31) were simulated; one in which all four consecutive carbon atoms were in trans conformation, and the other in which only one set of four consecutive carbon atoms was in gauche conformation (Fig. 2B). Simulated Raman spectra qualitatively reproduced the spectral transitions observed in Fig. 1, with increased width of the largest peak and a right shift of the peak top for 16:0(d31) with one gauche conformation (Fig. 2C, D and E). Similar spectral changes were confirmed for a majority of the identical simulations performed for C16:0(d31) with one gauche conformation at varying positions (Fig. S2). These results suggest that differences in gauche/trans conformations affect the spectral shape, confirming that C16:0(d31) is suitable as a probe to monitor the gauche/trans ratio of the lipid environment.

**Fig. 2.**
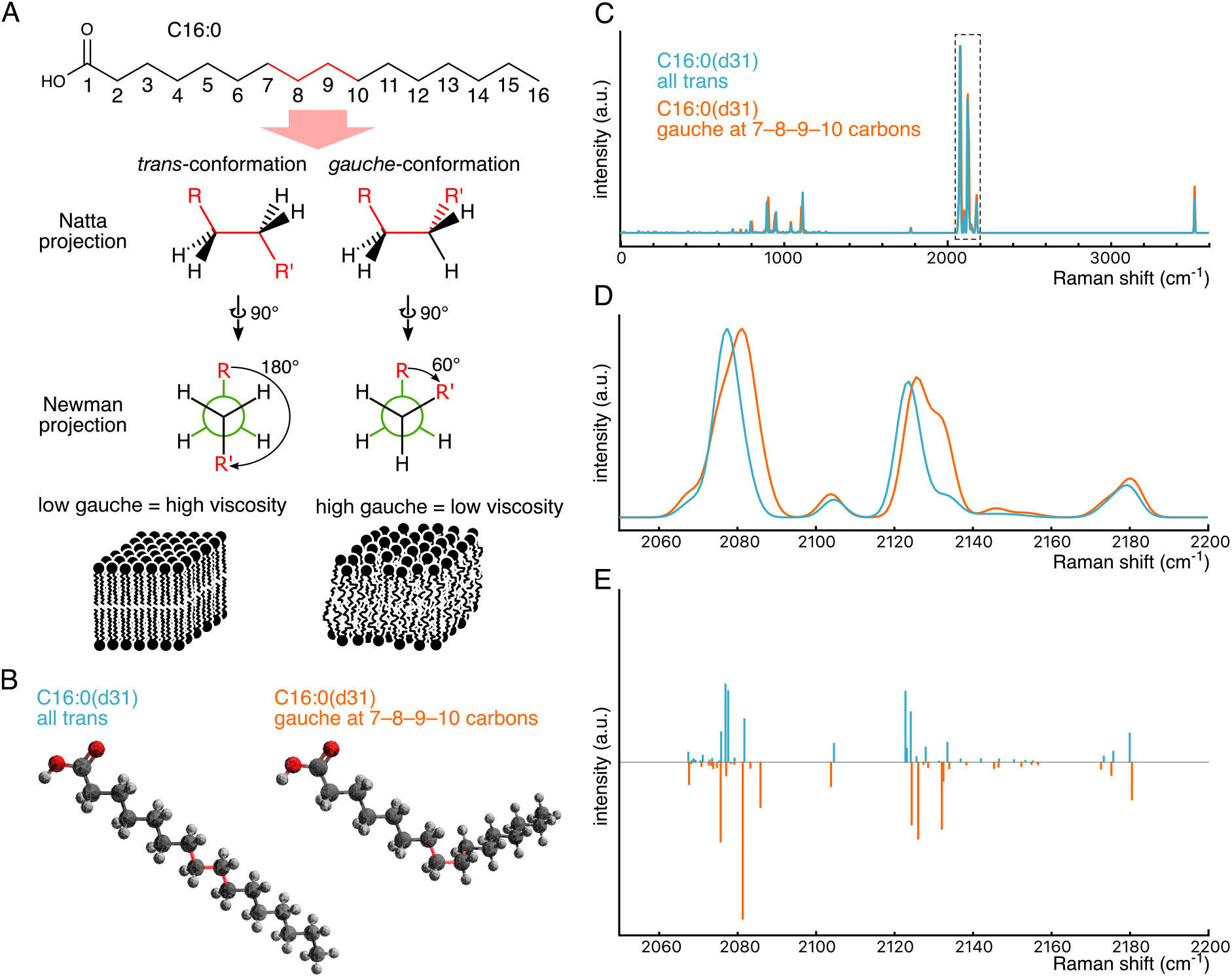
Gauche/trans conformation affects the spectra of C-D vibration. (**A**) Schematic images of gauche/trans conformation and membrane viscosity. Gauche/trans conformation is shown with Natta and Newman projections. Lipid bilayers with a lower (left) and higher (right) gauche/trans ratio have higher and lower viscosity, respectively. The numbering of carbon atoms in C16:0 is displayed in the skeletal formula. (**B**) Structures of C16:0(d31) with all trans (left) and one gauche conformation at 7-8-9-10 consecutive carbons (right). Dark gray, light gray, and red atoms represent carbon, deuterium, and oxygen atoms, respectively. Chemical bonds between four consecutive carbon atoms corresponding to the skeletal formula in (A) are shown with red. (**C**) Raman spectra simulated using DFT. Magnified view of the dashed box and its corresponding Raman activity are shown in (D) and (E), respectively. Intensity was normalized by equalizing prominent peak heights of the C–D vibration region. Graphs (D) and (E) are also shown in Fig. S2A and S2B.

Notably, although *in silico* analyses reproduced qualitative spectral transitions, they were unable to replicate the exact shape of experimentally observed spectral transitions, as shown in Fig. 1 and Fig. S4A. One possibility for this discrepancy is that one C16:0(d31) molecule contains thirteen positions that can possess a gauche/trans conformation, and that *in vitro* Raman spectra are derived from a mixture of heterogeneous conformational isomers containing various combinations of gauche/trans conformations at each position. Since our *in silico* simulation calculated the spectra of only one conformational isomer of one molecule at a time, it was challenging to reproduce the exact spectra of such a heterogeneous lipid environment.

### Spectra of C16:0(d31) are sufficiently sensitive to detect physical properties of different lipid environments *in vivo*

Thus far, *in vitro* and *in silico* analyses have demonstrated that C16:0(d31) is a promising probe for detecting the physical properties of lipid environments. Next, we assessed its application *in vivo* to observe the intracellular lipid environment under moderate physiological changes. Hence, we attempted to capture the *in vivo* spectral transitions of C16:0(d31) incorporated into HeLa cells associated with fatty acid composition and subcellular localization. HeLa cells were incubated with different compositions of fatty acids: C16:0(d31) mixed with C16:0(d0) or C20:4(d0) (30 μM each) (Fig. 3A) then, spectra in LD and non-LD regions were compared for each condition by subtracting the background spectra drawn with gray lines (see Materials and Methods). The results showed a rightward shift of the peak top of the spectrum in LDs of HeLa cells treated with the C16:0(d31)/C20:4(d0) mixture, compared to that with the C16:0(d31)/C16:0(d0) mixture, with the peak width widening (the top-right graph in Fig. 3A). These results suggest that the high degree of C20:4(d0) unsaturation resulted in a higher gauche/trans ratio of incorporated C16:0(d31) inside LDs. In contrast, such spectral transitions were hardly observed when comparing spectra of non-LD regions (the bottom-right graph in Fig. 3A). As for spectral differences between different subcellular locations, that is, LD and non-LD regions, the spectra of LDs showed a lower C16:0(d31) gauche/trans ratio than that of non-LD regions when treated with the C16:0(d31)/C16:0(d0) mixture (the inset in the bottom-left graph in Fig. 3A), while treatment with the C16:0(d31)/C20:4(d0) mixture resulted in a slightly higher gauche/trans ratio in the LD region (the inset in the bottom-middle graph in Fig. 3A).

**Fig. 3.**
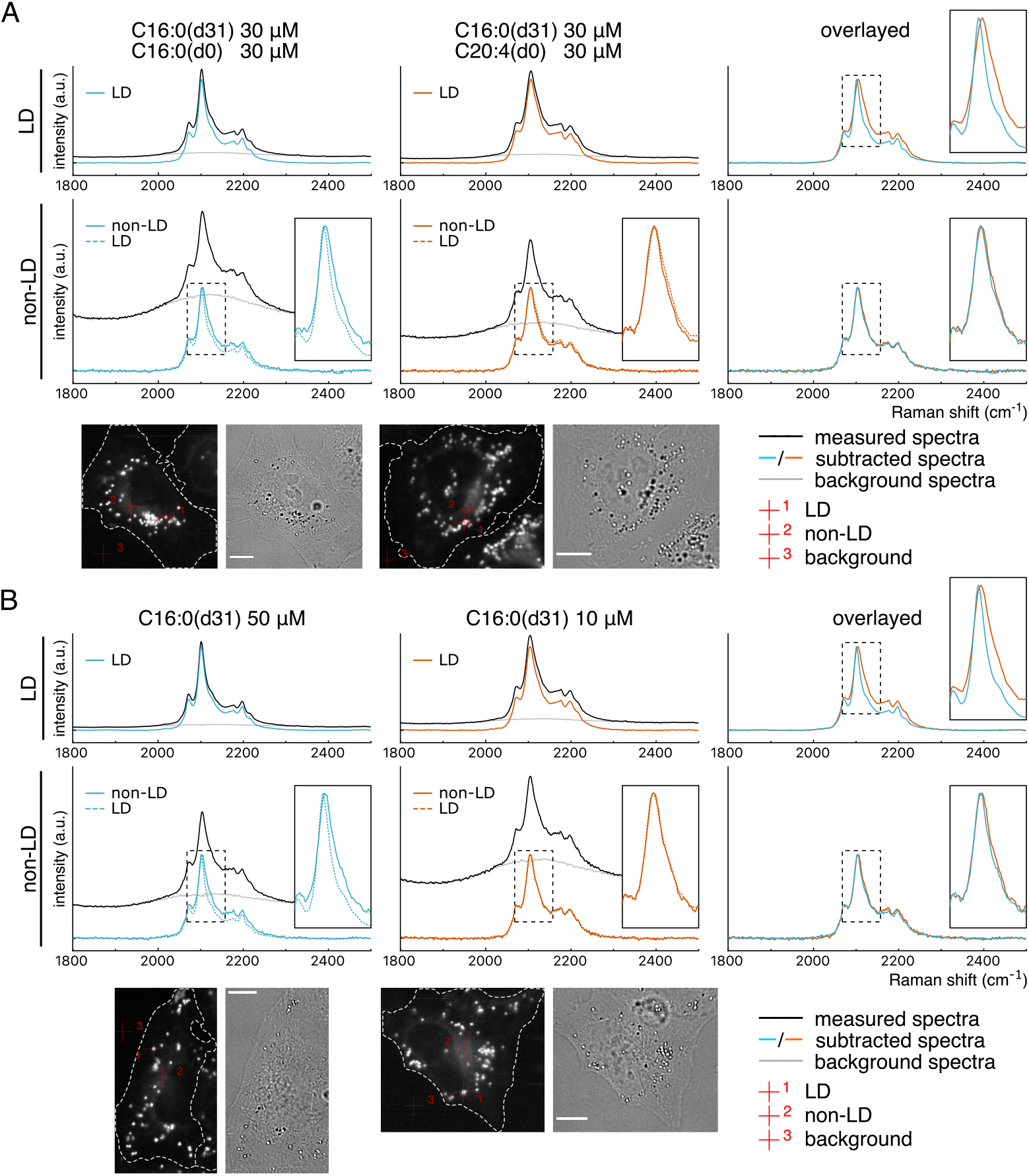
Spectral transitions of C16:0(d31) in HeLa cells. (**A, B**) HeLa cells were treated with indicated concentrations of fatty acids or a fatty acid mixture, and the representative Raman spectra from C–D vibration were compared at LD and non-LD regions. Background spectra (gray lines) were subtracted from each measured spectrum (black lines) with the appropriate weightings (see Materials and Methods) to illustrate a clear comparison, thus, producing subtracted spectra (solid blue or orange lines). Spectra from LD regions are also displayed with those from non-LD regions using dashed blue or orange lines at the bottom-left and bottom-middle graphs. Magnified views of the dashed black boxes are shown in the inset of some graphs. Pixels of representative LD regions, non-LD regions, and pixels used for background spectra are represented by red crosshairs 1, 2, and 3, respectively, in the bottom images. Two independent experiments confirmed the results. Scale bars indicate 10 μm.

Similar spectral transitions were also observed for HeLa cells treated with different concentrations of C16:0(d31) (Fig. 3B). Under the condition treated with lower concentrations (10 μM) of C16:0(d31), spectra derived from LDs were wider and had a rightward-shifted peak top compared to the spectra of LDs treated with higher concentration (50 μM) of C16:0(d31), suggesting a higher gauche/trans ratio (the top-right graph in Fig. 3B). In contrast, spectra of non-LD regions were relatively constant between the two conditions (the bottom-right graph in Fig. 3B). When spectra from LD and non-LD regions were compared in the same cells, non-LD regions showed a higher gauche/trans ratio than LDs in HeLa cells treated with 50 μM of C16:0(d31) (the inset in the bottom-left graph in Fig. 3B), while such spatial differences could hardly be observed in HeLa cells treated with 10 μM of C16:0(d31) (the inset in the bottom-middle graph in Fig. 3B).

These results provide methodological significance, as well as biological insight, into the function of LDs. The conformational state of C16:0(d31) in LDs was found to be diverse depending on the amount and composition of treated fatty acids; hence, it can be speculated that LDs are comprised of endogenous unsaturated fatty acids as well as excess C16:0(d31) taken up from the medium. Meanwhile, although the spectra of LD regions were highly dynamic in both Fig. 3A and Fig. 3B, those of non-LD regions were relatively constant, suggesting that the lipid environment in non-LD region is robustly preserved probably by LD serving as a buffer for excessive amounts of fatty acids inside cells.

### Generation of reference spectra to quantify spectral transitions

Although we have discussed the subcellular gauche/trans ratio in a qualitative manner based on the spectral transitions of C16:0(d31), a quantitative analysis is required for more accurate evaluation of the intracellular lipid environment. Thus, we next developed a system to quantitatively convert the spectral transitions of C16:0(d31) into gauche/trans ratio values. The quantification procedure comprises the following two steps: translation of the measured *in vivo* spectrum into the compositional information of the phospholipid mixture based on its *in vitro* reference spectra, and the calculation of the gauche/trans ratio of the *in vitro* phospholipid mixture using *in silico* model membrane simulations (Fig. 4A).

**Fig. 4.**
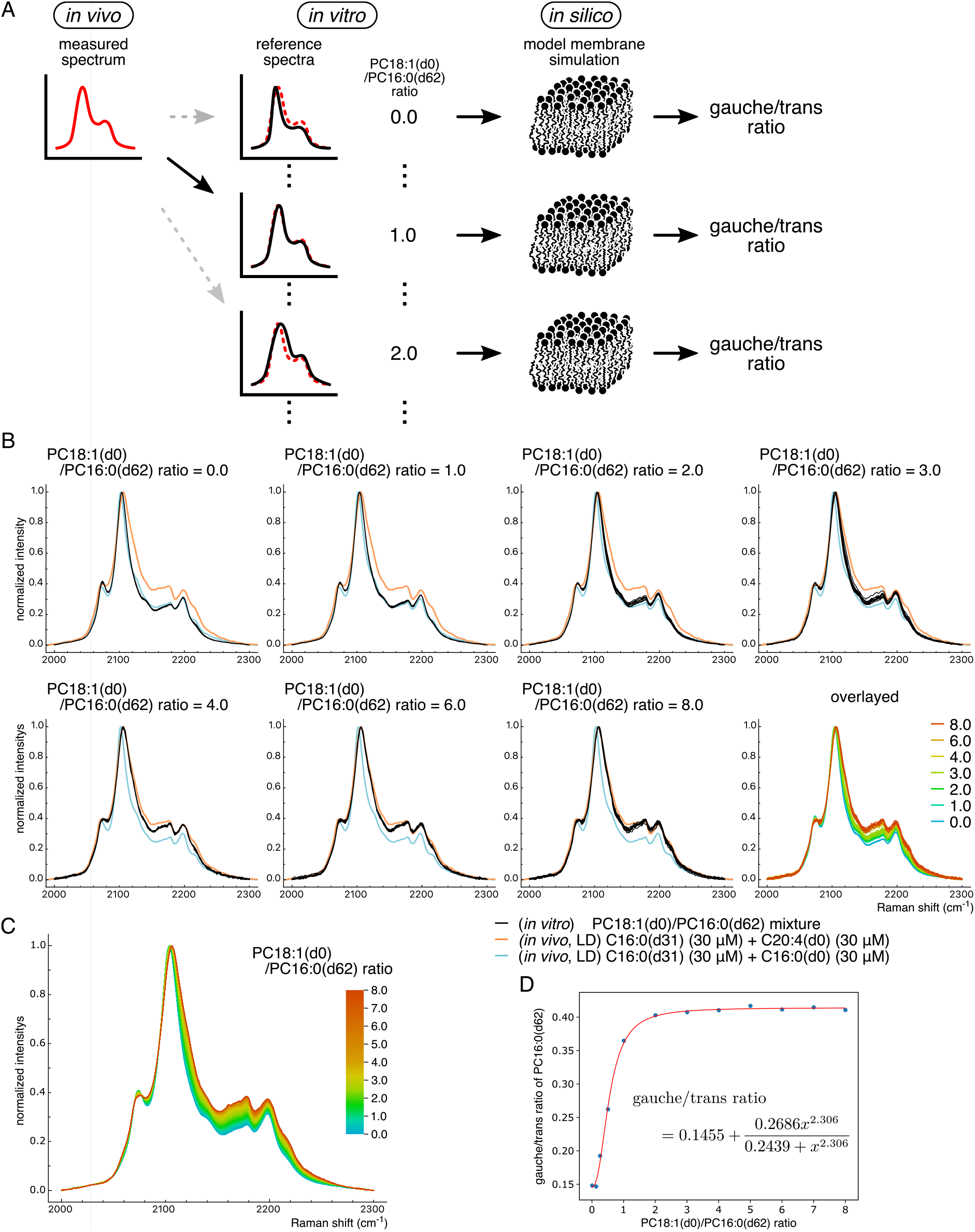
Quantification of measured spectra with the PC18:1(d0)/PC16:0(d62) and gauche/trans ratios. (**A**) Schematic image of the procedure for quantifying *in vivo* spectrum. Reference spectra of C–D stretch were first prepared from PC18:1(d0)/PC16:0(d62) phospholipid mixtures *in vitro*. Next, the PC18:1(d0)/PC16:0(d62) ratio that best explains the measured cell spectrum *in vivo* was estimated. The obtained ratio value was further converted into a gauche/trans ratio using the MD simulation of a model phospholipid membrane consisting of the same ratio of PC18:1(d0) to PC16:0(d62) *in silico*. Note that only discrete ratio values are shown in the scheme, but they were actually interpolated to yield continuous values. (**B**) *In vitro* spectra of PC18:1(d0)/PC16:0(d62) phospholipid mixture used to generate the reference spectra. Phospholipids were mixed in the indicated ratio, and the Raman spectra were measured. For each condition, spectra were measured from 9 or 10 different locations of the sample, and are displayed in the same graph using black lines (seven graphs excluding the bottom right). To show the similarity between *in vitro* and *in vivo* spectra, *in vivo* spectra of LDs from Fig. 3A are also displayed with blue and orange lines. The bottom right graph is an overlaid graph of the other seven graphs. (**C**) Reference spectra generated by the interpolation of *in vitro* spectra displayed in (B). Note that, color bar scale is not the same as in (B). (**D**) Relationship between the gauche/trans ratio and the PC18:1(d0)/PC16:0(d62) ratio estimated using *in silico* simulation. Results of the MD simulations are displayed as blue dots, and the red line indicates the function to express the gauche/trans ratio by the PC18:1(d0)/PC16:0(d62) ratio *x*. The function was obtained by the optimization of parameters based on the kinetic model (see Materials and Methods).

For the first step, we searched for a suitable combination of lipid species that could reproduce the *in vivo* spectral transitions observed in Fig. 3. In addition to the free form of C16:0(d31) used for treatment in the previous section, two types of lipids containing C16:0(d31) in their substructures were prepared and investigated: lysophosphatidylcholine (LPC) 16:0(d31), and phosphatidylcholine (PC) 16:0(d62) (Fig. S3). C16:0(d31), LPC16:0(d31), and PC16:0(d62) were mixed with C18:1(d0), C18:1(d0), and PC18:1(d0), respectively, at various ratios, and their Raman spectra were observed. The results showed that the PC16:0(d62) and PC18:1(d0) mixture could mimic the spectral transitions observed *in vivo* well (Fig. 4B), where the PC18:1(d0)/PC16:0(d62) ratio equaling 0 and 8 showed almost the same *in vitro* spectra as the LD spectra in HeLa cells treated with C16:0(d31)/C16:0(d0) (blue lines in Fig. 4B) and C16:0(d31)/C20:4(d0) mixtures (orange lines in Fig. 4B) from Fig. 3A, respectively. In addition, spectra gradually changed depending on the PC18:1(d0)/PC16:0(d62) ratio, with these intermediate spectra also reproducing the shape of *in vivo* spectra (data not shown). These spectral similarities indicate a high degree of resemblance in the conformational state of C16:0(d31) incorporated into cells *in vivo* and the fatty acyl chain of PC16:0(d62) *in vitro*; thus, these *in vitro* spectra from PC mixtures were chosen as the reference for evaluating *in vivo* spectra.

The mixture of free fatty acids C16:0(d31) and C18:1(d0) also showed spectral transitions (Fig. S4A). However, despite various adjustments of the mixing ratio of these fatty acids, the *in vivo* spectra of LDs in HeLa cells treated with the C16:0(d31)/C16:0(d0) mixture (blue lines in Fig. S4A) could not be reproduced. In addition, the reproducibility of *in vitro* spectra was poor when the C18:1(d0)/C16:0(d31) ratio was equal to 2, showing various patterns depending on the sample positions. These results indicate that C18:1(d0) and C16:0(d31) were not fully mixed. In contrast, the LPC16:0(d31) and C18:1(d0) mixture showed spectral changes similar to that of the PC mixture, also reproducing the *in vivo* spectra well (Fig. S4B). Nevertheless, considering that the LPC16:0(d31) spectra changed relatively steeply depending on the amount of C18:1(d0) (Fig. S4C), and that PCs are more abundant than LPCs in cells, spectra from PC mixtures were chosen as the reference.

Next, reference spectra were generated against continuously varying PC18:1(d0)/PC16:0(d62) ratios. Briefly, the raw spectra of phospholipid mixtures were measured with ratio ranges from 0 to 8 at finer intervals of 0.5, then obtained spectral data sets were interpolated. First, spectra intensities were normalized, and the intensities at each spectral data point along the wavenumber axis were plotted against the PC18:1(d0)/PC16:0(d62) ratio (Fig. S5). Next, piecewise cubic Hermite interpolation polynomial (PCHIP)-based regression was performed to obtain a continuous function from discrete data points, thereby generating spectra of the PC18:1(d0)/PC16:0(d62) mixture at an arbitrary ratio (Fig. 4C).

### Molecular dynamics (MD) simulation for the calculation of quantitative gauche/trans ratio values

For the second step of the quantification of the spectral transition, the gauche/trans ratio of acyl chains in PC16:0(d62) was calculated using MD simulations. Phospholipid model bilayer membranes with a PC18:1(d0)/PC16:0(d62) ratio of 0.0, 0.125, 0.25, 0.5, 1.0, 2.0, 3.0, 4.0, 5.0, 6.0, 7.0, or 8.0 were prepared *in silico*, and the MD simulations were performed. As expected, the gauche/trans ratio increased as the relative concentrations of PC18:1(d0) increased, eventually reaching just over 0.4 (Fig. 4D). Simulated data were fitted to an equation that was calculated based on a kinetic model (see Materials and Methods), thereby allowing us to convert the PC18:1(d0)/PC16:0(d62) ratio into a gauche/trans ratio (red line in Fig. 4D).

Using this two-step quantification approach, observed *in vivo* spectra can be converted into a PC18:1(d0)/PC16:0(d62) ratio, and subsequently into a gauche/trans ratio.

### Quantification of spectral changes *in vivo*

Next, we applied the developed quantification method to the *in vivo* data shown in Fig. 3. For the first step, spectral data at each pixel were converted into the PC18:1(d0)/PC16:0(d62) ratio values that account for the *in vivo* spectrum with the minimum residual sum of squares (RSS), considering background spectra and baseline drifts (see Materials and Methods for details) (Fig. 5A and B). The generated PC18:1(d0)/PC16:0(d62) ratio images showed relatively uniform intensity in LD and non-LD regions, thus, confirming that the pixels analyzed in Fig. 3 reflect the region-wide physical properties of lipids. Representative fitting results at the same pixels as in Fig. 3 were also shown. The sum of the estimated spectra (dashed blue and orange lines) fitted well with the measured spectra (solid black lines), suggesting that *in vitro* spectra are a good approximation of *in vivo* spectra. For the second step, PC18:1(d0)/PC16:0(d62) ratio images were converted into gauche/trans ratio images using the function obtained from the results of the MD simulations (Fig. 5C and D). The results also showed uniform gauche/trans ratios in LD and non-LD regions.

**Fig. 5.**
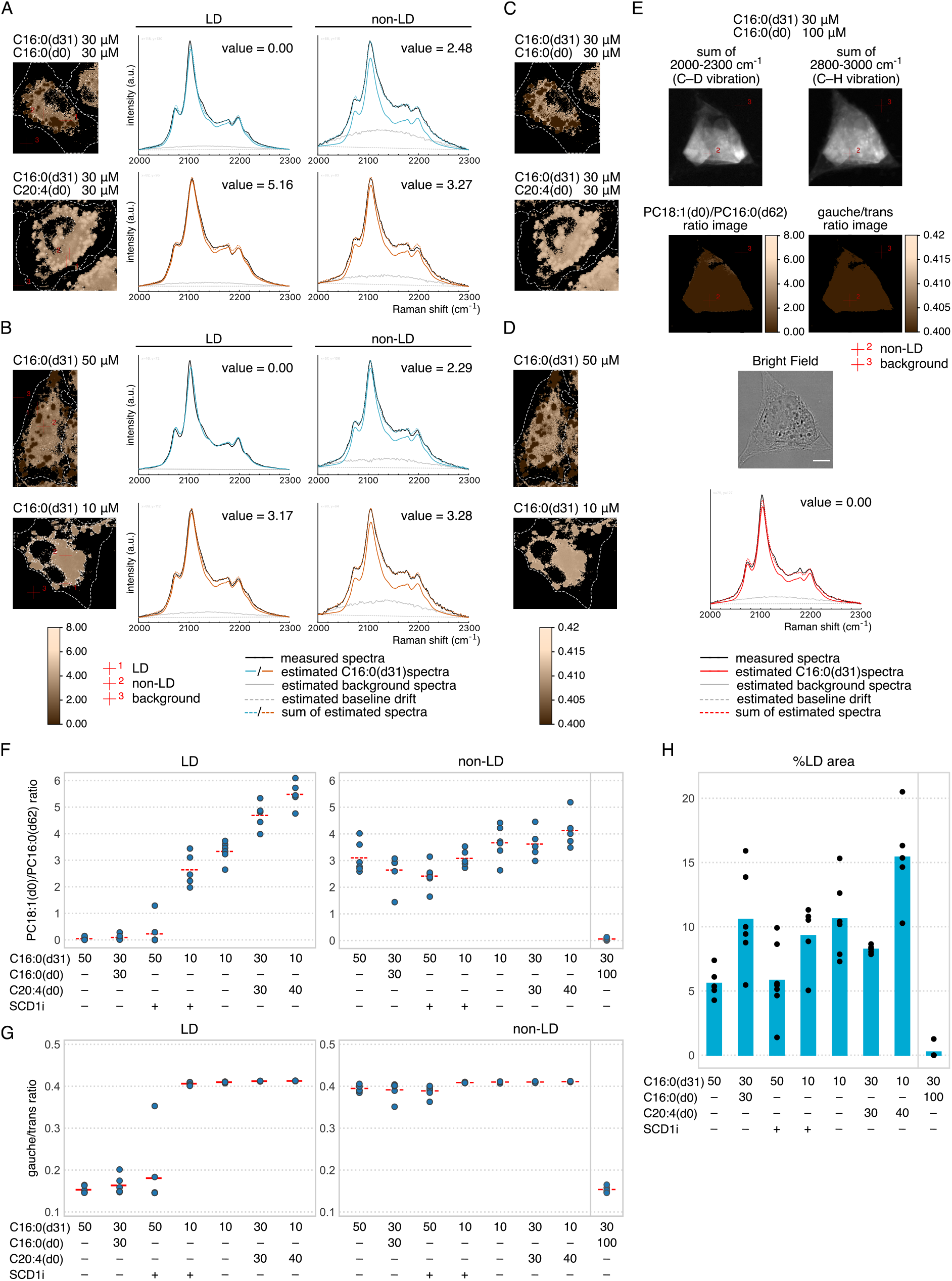
Quantification of *in vivo* spectra revealed the preserved physical properties in non-LD regions. (**A, B**) Raman hyperspectral data in Fig. 3 were converted into PC18:1(d0)/PC16:0(d62) ratio images. Representative results of the fitting at the same pixel as Fig. 3 (represented by red crosshairs 1 and 2) are displayed in the graphs on the right, with the calculated PC18:1(d0)/PC16:0(d62) ratio values. The background spectra from the same pixel as Fig. 3 (represented by red crosshair 3) were used for calculation. (**C, D**) PC18:1(d0)/PC16:0(d62) ratio images in (A) and (B) were further converted into gauche/trans ratio images. (**E**) HeLa cells treated with in total 130 μM of C16:0. Representative result of the fitting at the pixel represented by red crosshair 2 is displayed in the bottom graph, with the calculated PC18:1(d0)/PC16:0(d62) ratio value. The background spectra from the pixel represented by red crosshair 3 were used for calculation. The PC18:1(d0)/PC16:0(d62) and gauche/trans ratio images are displayed with the same color scaling as (A) to (D). The C–D and C–H vibration images, and the bright field image are displayed to show the distribution patterns of C16:0(d31) and the shape of the cell. In the graph of (A), (B), and (E), measured spectra are indicated with black lines, and estimated C16:0(d31) spectra, background spectra, baseline drift, and their summation are displayed with solid-colored lines, solid gray lines, dashed gray straight lines, and dashed colored lines, respectively. (**F, G**) Comparison of quantified PC18:1(d0)/PC16:0(d62) and gauche/trans ratios between LD and non-LD regions. Average ratio of LD and non-LD regions were calculated for five to seven cells and plotted. Red horizontal dashed lines indicate the average value for each condition. Conditions are ordered according to the average PC18:1(d0)/PC16:0(d62) ratio value in the LD region. (**H**) Quantification of the ratio of LD area to the sum of LD and non-LD area. Ratio values were calculated using the same data set as (F) and (G), and average values and each data point are represented with blue bar-graphs and black dots, respectively. Scale bar indicates 10 μm.

The trends of the physical properties in LD and non-LD regions were further analyzed by calculating the average PC18:1(d0)/PC16:0(d62) and gauche/trans ratio of each region over multiple cells. In addition to the four conditions described in Fig. 5A, B, C, and D, the following conditions were introduced to the analysis: treatment with the fatty acid mixture with a higher proportion of C20:4(d0), and pharmacological treatment using an inhibitor of stearoyl-CoA desaturase 1 (SCD1) (Fig. 5F and G, except for the rightmost condition in the non-LD region). Note that, SCD1 is an enzyme which introduces a double bond to the carbon at position 9 of C18:0-CoA and C16:0-CoA to produce C18:1-CoA and C16:1-CoA, respectively^17^. For the first condition, HeLa cells were treated with a C16:0(d31)/C20:4(d0) mixture (10 μM/40 μM); for the second condition, cells were treated with a mixture of 10 or 50 μM of C16:0(d31) and a SCD1 inhibitor. As expected, PC18:1(d0)/PC16:0(d62) and gauche/trans ratios were the highest among all conditions in HeLa cells treated with C16:0(d31)/C20:4(d0) (10 μM/40 μM) for both LD and non-LD regions. As for SCD1 inhibitor treatment, PC18:1(d0)/PC16:0(d62) ratios slightly decreased in LD and non-LD regions of HeLa cells supplemented with 10 μM of C16:0(d31) and in non-LD regions of HeLa cells supplemented with 50 μM of C16:0(d31), compared to in the absence of SCD1 inhibitor. These results quantitatively detected the effect of SCD1 inhibition on the subcellular lipid environment, indicating that non-LD regions of HeLa cells treated with 10 μM of C16:0(d31) containing a SCD1 inhibitor were affected to the same extent as those treated with 50 μM of C16:0(d31) in the absence of a SCD1 inhibitor.

Notably, the PC18:1(d0)/PC16:0(d62) and gauche/trans ratios of LDs changed dramatically in these seven conditions, with average ratio values ranging from 0.045 to 5.48 and 0.153 to 0.413, respectively, while lipid environments in non-LD regions were relatively conserved, with ratio values ranging from 2.42 to 4.12 and 0.390 to 0.411, respectively. These results suggest the possible mechanism that keeps the physical properties of cytoplasmic lipid environment stable, probably by LDs acting as buffers for excessive degrees of the gauche/trans ratio. To explore the contribution of LDs to this robust lipid environment in non-LD regions, we searched the conditions under which this robustness is lost. As a result, we found that the treatment with higher amount of C16:0 (130 μM in total) greatly reduced the PC18:1(d0)/PC16:0(d62) and gauche/trans ratios in non-LD regions (Fig. 5E, F and G). Meanwhile, almost no LD formation was observed under this condition (Fig. 5H), and cell viability was greatly reduced at the same time. These results strongly suggest that the normal function of LDs to accumulate excess fatty acids is crucial for the maintenance of the lipid environment in non-LD regions, and consequently is necessary for cell survival.

## DISCUSSION

In the present study, we developed a Raman microscopy-based method for monitoring the subcellular physical properties of a lipid environment as a gauche/trans ratio. Gauche/trans ratios and lipid viscosity are intimately related with one another, with their correlations strongly suggested through experiments manipulating viscosity at different temperatures^13,16^. Thus, although a direct relationship was not demonstrated in this study, our method may be useful to indicate the viscosity of a lipid environment.

A fundamental difference between our approach and previous techniques is that our method uses the lipid itself as a probe, thus enabling us to directly measure the physical properties of lipids without causing disruption by the introduction of probes. Although our method is limited to fatty acid-treated conditions, such perturbations are commonly used to study intracellular lipid transportation and LD functions, and should therefore allow us to study these processes under more natural conditions compared to other existing techniques. In addition, the increase of the sensitivity of Raman microscopes in the future will reduce the amount of C16:0(d31) required to be introduced, thereby making it possible to measure the gauche/trans ratio of subcellular lipid environments under a wider range of conditions.

A Raman microscopy-based approach different from ours was recently reported to monitor the physical properties of lipids as a lateral diffusion coefficient^12^. The diffusion coefficient is commonly used to represent the physical properties of lipids, and is known to be directly related to membrane viscosity. They estimated this value by observing the ripple-like patterns formed by pulse-labeling with C16:0(d31) for one hour. In this sense, these newly discovered domains are appropriate targets for monitoring the lipid environment, as they are relatively large (around 10 μm) structures that can grow stably for several hours while continuously taking up C16:0(d31). In other words, it is unclear whether this method can be applied to other much smaller and more highly dynamic organelles. They also performed a ratio metric analysis utilizing the transition of C16:0(d31) spectra to confirm that domains were actually phase-separated. Although this method can partially quantify the lipid environment, calculating the ratio of just two spectral data points is unstable, especially where intensity of the C16:0(d31) signal is weak and the effect of the background signal is ignorable. In addition, it was ambiguous what aspects of the lipid environment the spectral transition represented. In contrast, our approach compensates for these issues, revealing the spectral transition is reflecting the gauche/trans ratio of incorporated C16:0(d31) at the same time.

Accumulation of C16:0(d31) in LDs enabled us to investigate and identify new biological insights into the physical properties of LDs, which has also been difficult to perform using conventional methods due to the lack of appropriate probes targeting LDs. The gauche/trans ratio of LDs changed dramatically in response to fatty acid treatment or pharmacological intervention, while that of non-LD regions remained relatively constant (Fig. 5F and G). These results imply the existence of some mechanisms through which the cytoplasmic lipid environment is maintained, possibly with LDs acting as a buffer for excessive “gauche/trans ratio”. Meanwhile, such robustness of non-LD regions was lost when LD formation was not observed by the treatment with high concentration of C16:0 (130 μM in tltal) (Fig. 5E, F, G and H). Although localization patterns of incorporated labeled fatty acids in non-LD regions are unclear, previous reports including ours suggest they mainly localize to ER, where LDs are generated^12,14^. Considering that decreased unsaturation of phospholipids has been reported to antagonize the formation of LDs and lipoproteins^5,18^, the loss of this robustness in non-LD region could be due to falling into a negative spiral of decreased unsaturation in ER, initiated by the treatment with high doses of C16:0 beyond the capacity of the buffering function of LDs to maintain the gauche/trans ratio of ER and its function to generate LDs. Biological phenomena we found based on the visualization of the subcellular physical properties of lipids offer new insight into the functions of LDs, which were previously regarded as only an energy storage organelle.

ER is an important organelle involved in the synthesis, folding, and transport of a majority of proteins. Abnormal increases in saturated fatty acids or decreases in unsaturated fatty acids is known to trigger ER stress via an unfolded protein response (UPR), subsequently leading to cell apoptosis^19^. On the other hand, increased unsaturated fatty acids in the phospholipid membrane also induce cytotoxicity in the similar manner via UPR-mediated apoptosis^20^. These reports demonstrated that imbalance of saturated and unsaturated fatty acids leads to ER stress, and the physical property of ER membranes should be vital to maintain the normal ER functions. LDs may serve as a buffer for this, and our results that gauche/trans ratio in non-LD region is constantly preserved support this concept. Since ER is involved in a wide range of cellular functions, our approach could be useful for revealing the more general mechanisms of cellular robustness to external stimuli through the physical properties of ER membranes.

### Limitations

There are some limitations to our methods; the assumption that the gauche/trans ratio is a dominant modulator of the spectral transitions of C–D stretch, and insufficient dynamic ranges.

For the first point, spectra derived from C–D stretch were converted into the PC18:1(d0)/PC16:0(d62) ratio, and further into the gauche/trans ratio during the quantification processes. In the process of converting the PC18:1(d0)/PC16:0(d62) ratio into the gauche/trans ratio, we used DFT to demonstrate that the spectral transitions were generated by changes in the gauche/trans ratio, and that the gauche/trans ratio were actually changed by the PC18:1(d0)/PC16:0(d62) ratio of model membranes in the MD simulation. However, it cannot be ruled out that similar spectral transitions may be induced by other factors of molecular states. For example, inter-molecular interactions may also affect the vibrational state of the C–D stretch. In fact, in the spectra of water molecules, strength of the hydrogen bonding is known to affect the shape of Raman spectrum derived from O–H stretch. Since the density of molecules dissolved in water affects the strength of hydrogen bonding, this phenomenon is used to measure subcellular water concentration^21^. Further studies, both *in vitro* and *in silico,* are required to more accurately estimate the physical properties of PC18:1(d0)/PC16:0(d62) phospholipid mixtures.

The second limitation to our method is the insufficient dynamic range of quantification. When comparing the fitting of spectra from LD and non-LD regions of cells treated with a total of 60 μM of C16:0 (the top two graphs in Fig. 5A), spectra fittings from LDs were slightly worse than those from the non-LD regions. The peak top of measured spectra from the LD region (black line in the top left graph of Fig. 5A) was located more to the left than the best-fitted spectra (dashed blue line in the top left graph of Fig. 5A), suggesting that the gauche/trans ratio of LDs in this condition was lower than that of phospholipid mixture with a PC18:1(d0)/PC16:0(d62) ratio equal to 0. This may be due to the fact that LDs predominantly consists of triglycerides and cholesterol esters, allowing for them to be packed at a higher density than lipid films composed of only phospholipids. In fact, introducing cholesterols is known to reduce the fluidity of phospholipid membranes, by helping phospholipid molecules being packed with higher density. We used a pure phospholipid mixture model as the *in vitro* reference sample this time, but introducing triglycerides or cholesterols to the model would enable the measurement of the physical properties of lipids with higher dynamic ranges.

## MATERIALS AND METHODS

### Labeled and non-labeled lipids

C16:0(d2) (48966-2) and C18:0(d3) (49039-3) were purchased from Taiyo Nippon Sanso (Tokyo, Japan). C18:0(d5) (D-5400) was purchased from C/D/N Isotopes (Tokyo, Japan). C20:0(d4) (DLM-10519-PK) was purchased from Cambridge Isotope Laboratories (MA, USA). C16:0(d31) (16497), C18:1(d17) (9000432), C18:2(d4) (390150), C18:2(d11) (9002193), C18:3(d14) (9000433), C20:4(d8) (390010), C20:4(d11) (10006758), C20:5(d5) (10005056), C22:6(d5) (10005057), C16:0(d0) (10006627), C18:1(d0) (90260), and C20:4(d0) (90010) were purchased from Cayman Chemical (MI, USA). PC16:0(d62) (860355), PC18:1(d0) (850375), and LPC16:0(d31) (860397) were purchased from Avanti Polar Lipids (AL, USA). For C16:0(Br) and C18:0(Br), chemically synthesized compounds in a previous study were used^15^.

### Cell culture

HeLa cells were cultured in Dulbecco’s modified Eagle’s medium (08459; Nacalai Tesque, Kyoto, Japan) supplemented with 10% fetal bovine serum (12676-029; Thermo Fisher Scientific, Waltham, MA, USA), in a 5% CO_2_ incubator. To conduct imaging experiments using labeled free fatty acids, HeLa cells were seeded on 35-mm quartz bottom dishes (SF-S-D12; Techno Alpha, Tokyo, Japan). After 24 h of incubation, the media were replaced with the desired combination and concentration of labeled and/or non-labeled free fatty acid solutions, and further incubated for 24 h. Cells were washed with phosphate-buffered saline (PBS), fixed with 4% paraformaldehyde (160-16061; Wako, Osaka, Japan) for 20 min at room temperature, and then again washed five times with PBS. To conduct the SCD1 inhibition experiment, HeLa cells were treated with a SCD1 inhibitor (ab142089; Abcam, Cambridge, United Kingdom) 1 h before free fatty acid treatment.

### Raman imaging

To obtain Raman hyperspectral images of cells, samples were set onto a confocal Raman microscope (inVia Reflex; Renishaw, Wotton-under-Edge, United Kingdom), and excited using a 532-nm diode laser through a 63× water immersion objective (506148, NA = 0.90; Leica, Wetzlar, Germany), followed by separation of scattered light using either 600 or 1800 line/mm grating. Exposure time and stage movement intervals were set to 0.1 s/pixel and 0.3 μm/pixel, respectively, using WiRE 5.1 software (Renishaw).

### Raman spectroscopy

To obtain the Raman spectra of compounds, free fatty acids, LPC, and PCs were dissolved in ethanol, a mixed solution of water/ethanol (1:1 v/v), and chloroform, respectively. Next, aliquots (0.5 μL) of each solution, or a mixture of them, were dropped onto CaF2 glass substrates (OPCFU-25C02-P; Sigma Koki, Tokyo, Japan).Following evaporation of the solvent, Raman spectra were obtained using WiRE 5.1 software and confocal Raman spectroscopy at room temperature. Samples were excited using a 532-nm diode laser through a 50× air objective (566072, NA = 0.75; Leica) followed by separation of scattered light using either 600 or 1800 line/mm grating.

### *In silico* simulation of Raman spectra

Design and geometry optimization were performed using Avogadro version 1.2.0^22^, based on molecular mechanics with a universal force field. Next, quantum chemical calculations were performed to infer Raman spectra using DFT at the B3LYP level with 6-31pG(d) basis set in Orca version 4.2.1^23,24^. For visualization, a linear scaling factor of 0.945 and a Gaussian width of 3 were applied.

### Calculation of the reference spectra by the interpolation of Raman spectra of the PC18:1(d0)/PC16:0(d62) mixture

Intensity of the reference spectra at the *s*-th data point along the wavenumber axis was obtained as a function of the PC18:1(d0)/PC16:0(d62) *x*, which is represented as *f*_*s*_ (*x*) in the following explanation, by PCHIP-based regression. Phospholipid mixtures with the PC18:1(d0)/PC16:0(d62) ratio varying from 0 to 8 in increments of 0.5 were prepared, and their Raman spectra were measured. For each condition, 9 or 10 spectra from different positions of the sample were collected. Spectral regions ranging from 2000 to 2300 cm^−1^ (210 data points with approximately 1.43 cm^−1^ interval) were normalized after subtracting slope and then used for the following analysis. Then, the whole data set ***U*** can be represented as follows:

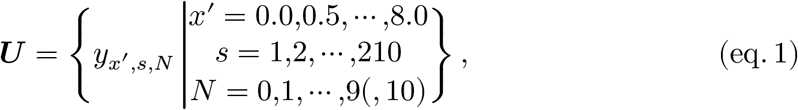

where *y*_*x*_′,*s*,*N* represents the intensity of the normalized spectrum of the phospholipid mixture at the *s*-th data point along the wavenumber axis when the ratio of PC18:1(d0)/PC16:0(d31) and the sample number equal *x*′ and *N*, respectively. Next, PCHIP-based regression was performed against sub-data set ***U***_*s*_′ to generate a piecewise cubic function *f*_*s*_′ (*x*) for each *s*′ using the PchipInterpolator class of SciPy 1.2.1^25^ and Python (version 3.6, https://www.python.org), where

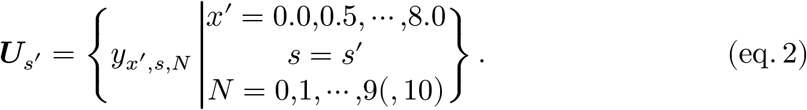

To determine *f*_*s*_′ (*x*), first, it was given the following two conditions. One is to pass through five coordinates 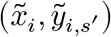 called “knots” for *i* = 0,1,2,3,4 where

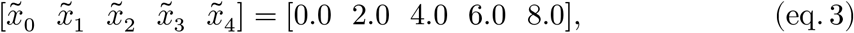

and 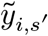 are five parameters to be optimized as discussed later in this section. The other condition is the differential coefficient of *f*_*s*_′ (*x*) at each knot, which is represented as 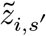 and is defined using temporal variables for 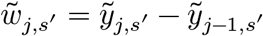 for *j* = 1,2,3,4, as follows:

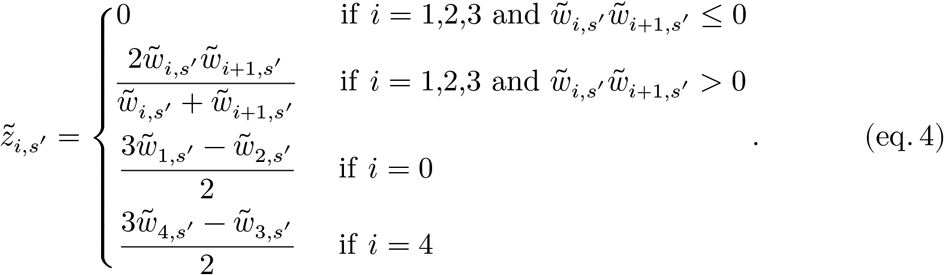

With these restrictions, *f*_*s*_′ (*x*) can be represented as a piecewise cubic function as follows:

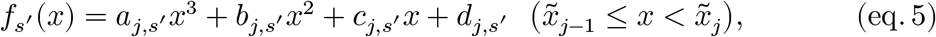

where

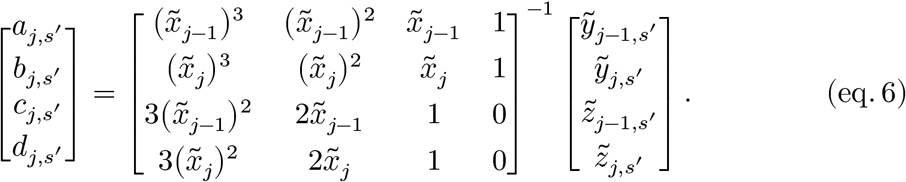

Then, parameters 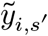 were optimized against *y*_*x*_′,*s*′,*N* ∈ ***U***_*s*_′ to minimize the following residual sum of squares (RSS_1_):

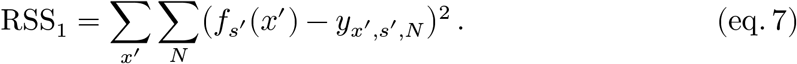

Finally, by using *f*_*s*_′ (*x*) with the optimized coefficients 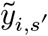, vector ***y**_x_*, which represents the reference Raman spectra of the phospholipid mixture with the PC18:1(d0)/PC16:0(d31) ratio equaling *x*, were generated in the range from 2000 to 2300 cm^−1^ as follows:

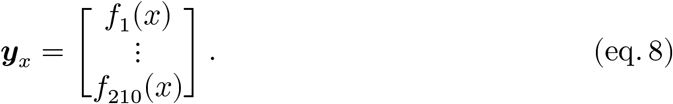

### Conversion of Raman spectra into the PC18:1(d0)/PC16:0(d62) ratio

The region between 2000 and 2300 cm^−1^ of the measured Raman spectra at each pixel (represented by vector *m*) was converted into virtual PC18:1(d0)/PC16:0(d62) ratio *x* by optimizing five parameters, including *x*, (*x*, *α*, *β*, *γ*, and *δ*), to minimize the following RSS_2_ with constraints of *α,β* ≥ 0:

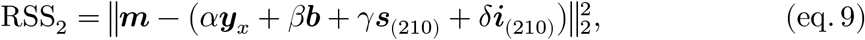

Here, ‖***a***‖_2_ represents the L2 norm of vector ***a***, and ***b*** represents the region between 2000 and 2300 cm^−1^ of the background spectrum. ***s***_(210)_ and ***i***_(210)_, corresponding to the slope and the intercept, respectively, are vectors with the same size as ***y***_*x*_ (which equals 210), and are represented as follows:

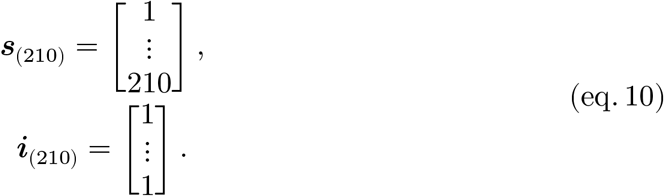

Calculations were performed using the minimize function of SciPy 1.2.1 and Python 3.6 implemented in ImageCUBE version 0.6.4^14^.

### Background subtraction of *in vivo* spectra

To simplify the comparison of the spectra, background spectra were subtracted with weighting from representative spectra of LD and non-LD regions. The weighting values *β*_LD_, *β*_nonLD_, *γ*_LD_, *γ*_nonLD_, *δ*_LD_, *δ*_nonLD_, were calculated to minimize the following RSS values:

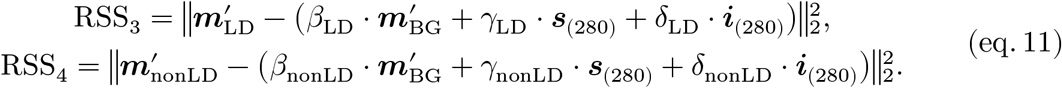

Here, 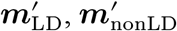, and 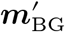 represent the concatenated two sub-vectors of the spectra measured in representative LD, non-LD, and background pixels, respectively. The size of each sub-vector was 140, and the region from which sub-vectors were extracted was 1800– 2000 and 2300–2500 cm^−1^. Of note, the procedure for this background subtraction was used only for the visualization of spectra in Fig. 3, and not for other analyses.

### MD simulation

Model membrane systems were created using CHARMM-GUI^26,27^ with the following PC18:1(d0)/PC16:0(d62) ratio (numbers in brackets followed by the ratio represent the numbers of each phospholipid molecule): 0.0 [0/200], 1.0 [100/100], 2.0 [136/68], 3.0 [150/50], 4.0 [160/40], 5.0 [170/34], 6.0 [180/30], 7.0 [182/26], and 8.0 [192/24]. A height of 22.5 nm was applied to the top and bottom of the membranes and filled with a TIP3P water model. Next, simulations were performed using the GROMACS 2019.3 software package^28^, with CHARMM36m all-atom force field^29^. First, system energy was minimized using the steepest descent method, followed by equilibration and production steps. In the equilibration steps, the following conditions were first applied for 250 ps: NVT ensemble with an integration time step of 1 fs, and a constant temperature of 298.15 K using the Berendsen thermostat^30^. In this step, position restraints of 1000 and 400 kJ mol^−1^ nm^−2^ were applied every 125 ps to the phosphorus atoms of each lipid. Additionally, dihedral restraints of 1000 and 400 kJ mol^−1^ rad^−2^ were applied to dihedral angles, defined by the following two types of four atoms (position of atoms are indicated in brackets after the atomic symbol): C[sn-1], C[sn-3], C[sn-2], O[sn-2] (for both PC16:0[d62], and PC18:1[d0]), and C[8], C[9], C[10], and C[11] of each acyl chain (for PC 18:1[d0]). Next, the system was further equilibrated in NPT ensemble equilibration for 875 ps using the Berendsen barostat with a constant pressure of 1 bar^30^. In this step, positional restraints of 400, 200, 40, and 0 (no restraints) kJ mol^−1^ nm^−2^, dihedral restraints of 200, 200, 100, and 0 kJ mol^−1^ rad^−2^, and time step of 1, 2, 2, and 2 fs were applied for the first 125 ps, and every subsequent 250 ps.

During production simulations, no restraints were applied to the systems, and the temperature and pressure were kept constant at 298.15 K and 1 bar, respectively, using Nosé–Hoover thermostat^31,32^ and a Parrinello-Rahman barostat^33^ with semi-isotropic pressure coupling to characteristic times of 1 and 5 ps, respectively. Throughout the simulation, all non-bonded interactions were cutoff at 1.2 nm.

### Trajectory analysis

We used GROMACS 2019.3, Python 3.6, gnuplot (version 5.2, http://www.gnuplot.info), and PyMol (version 2.3.4, https://pymol.org/2/) for the analysis and visualization of trajectories.

### Modeling of the gauche/trans ratio of the PC18:1(d0)/PC16:0(d62) mixture

Interconversion between gauche and trans conformers were represented schematically as

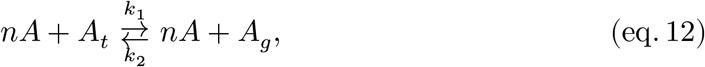

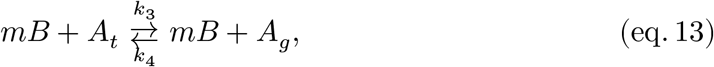

where *A*, *B*, *A_g_*, *A_t_*, *n*, *m* and *k_i_* (*i* = 1,2,3,4) denote the PC16:0(d62), PC18:1(d0), dihedral carbon atoms in PC16:0(d62) with gauche conformation, dihedral carbon atoms in PC16:0(d62) with trans conformation, order of reaction of eq.12, order of reaction of eq.13, and reaction rate constants, respectively. Based on these reaction formulae, the following equation can be established under equilibrium:

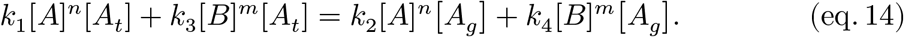

This equation can be transformed as follows to calculate the gauche/trans ratio:

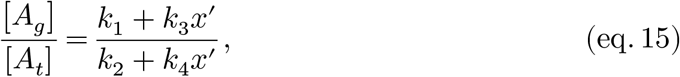

where

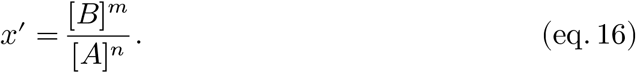

Assuming that the order of reaction was the same for eq.12 and eq.13 (*n* = *m*), the following equation was derived;

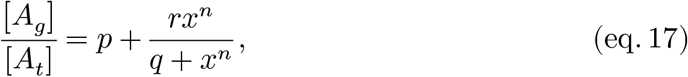

where

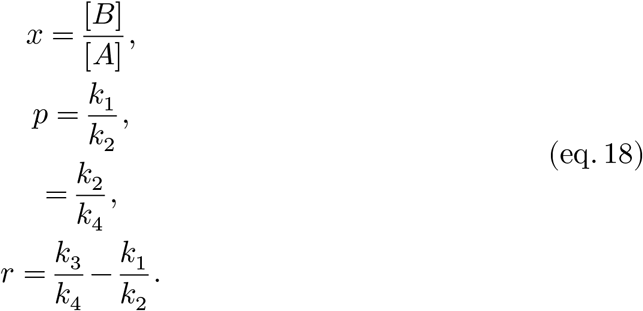

Thus, gauche/trans ratios were expressed with the PC18:1(d0)/PC16:0(d62) ratio *x* Parameters *p*, *q*, and *r* were optimized to fit the data derived from MD simulations using the minimize function of SciPy 1.2.1 and Python 3.6.

### Quantification of cell images

All images were analyzed using ImageCUBE version 0.6.4 and Fiji/ImageJ^34,35^. At the first quantification step, images were masked based on the signal-to-noise ratio (SNR_2_) and horizontal line artifacts (HLAs) deriving from microscopic scans. For each pixel, SNR_2_ was calculated as follows:

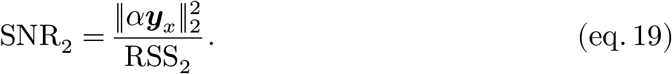

Here, *α*, ***y***_*x*_, and RSS_2_ are derived from eq.8 and eq.9. Images constructed using the value 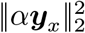, RSS_2_ and SNR_2_ are shown as 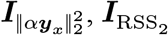, and 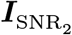, respectively, in Fig. S6. HLAs were defined for each image 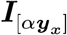 constructed based on the sum of the elements of *α****y***_*x*_ represented [*α****y***_*x*_] as follows:

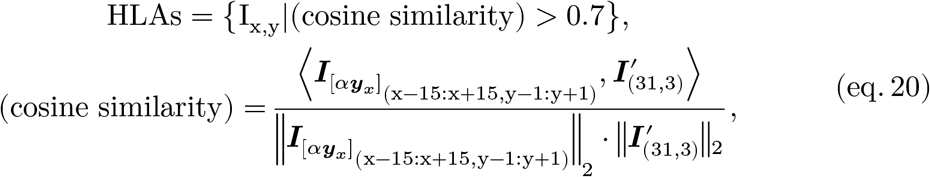

where **I**_x,y_ represents the pixel at (x, y), and the operations 〈***A,B***〉, and ‖***A***‖_2_ represent the inner product of matrices ***A*** and ***B*** and the L2 norm of matrix ***A***, respectively. 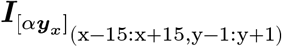 represents the matrix of sub-image 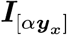, with x and y coordinates ranging from x − 15 to x + 15 and from y − 1 to y − 1, respectively. 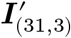 is a matrix of the same size as 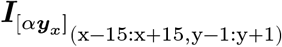 and is described as follows:

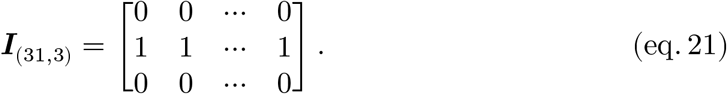

Next, HLAs were horizontally extended for 10 pixels to acquire HLAs_extended_. Intermediate images of the process are shown in Fig. S6 as ***I***_cos_, ***I***_HLAs_ and ***I***_HLAsEx_. Finally, pixels that satisfy SNR_2_ > 100, and not being included in HLAs_extended_, were subsequently analyzed (***I***_MASK_ in Fig. S6).

As a second step, LD and non-LD areas were defined from image 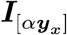. First, the background image ***I***_BG_ was generated using a median filter with a kernel size of 21, which was subsequently subtracted from image 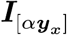. Next, image convolution was performed (***I***_CONV_ in Fig. S6) to detect pixels in which the brightness of the image drastically changed, using the following kernel ***K*** with size 5:

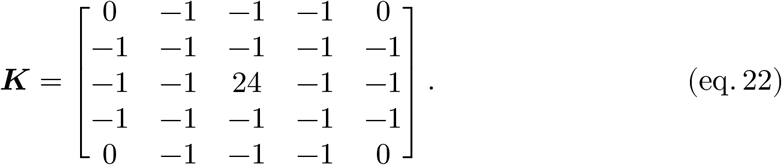

After convolution, the Fourier transformation (FT) bandpass filter was applied, filtering large and small structures down to 5 and up to 2 pixels, respectively(***I***_FT_ in Fig. S6). Next, the *threshold* was set as follows using the modal intensity of ***I***_FT_:

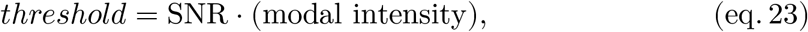

where SNR was arbitrarily set to 5. Finally, pixels whose intensities were greater than the *threshold* and that were included in ***I***_MASK_ were defined as LD areas (***I***_LD_ in Fig. S6). To define non-LD areas (***I***_nonLD_ in Fig. S6), LD areas were swelled by 2 pixels (***I***_sLD_ in Fig. S6) and subtracted from ***I***_MASK_.

Note that, as this procedure produces unexpected results when applied to cells without LDs, such cells were analyzed manually.

## ABBREVIATIONS

Br: bromine
DFT: density functional theory
ER: endoplasmic reticulum
FT: Fourier transformation
HLA: horizontal line artifact
LD: lipid droplet
LPC: lysophosphatidylcholine
MD simulation: molecular dynamics simulation
PBS: phosphate-buffered saline
PC: phosphatidylcholine
PCHIP: piecewise cubic Hermite interpolation polynomial
RSS: residual sum of squares
SCD1: stearoyl-CoA desaturase 1
SNR: signal-to-noise ratio
SRS: stimulated Raman scattering
UPR: unfolded protein response

## Code availability

The Python, ImageCUBE, and ImageJ code is available from the corresponding author upon request.

## Data availability

The data that support the findings of this study are available from the corresponding author upon request.

## ACKNOWLEDGMENT

We would like to express our gratitude to Hideo Shindou for providing a good research environment and for thorough discussion. We also thank Hiroshi Noguchi for providing valuable advice on MD simulation, Saori Uematsu, Keisuke Yanagida and Daisuke Hishikawa for their thoughtful discussions and review of the manuscript, Shota Yamamoto, Tomoyuki Suzuki, Natsuko Inagaki, Miyuki Yamamoto, Yukiko Sugimoto, and Megumi Yasuda for kindly allowing us to use their computers for the analyses. All MD simulation results in this paper were generated on the NIG supercomputer at ROIS National Institute of Genetics.

## AUTHOR CONTRIBUTIONS

M. Uematsu conceptualized the project, developed the analytical methods, performed the *in vivo*, *in vitro*, and *in silico* experiments, investigated the data, and wrote the manuscript. T. Shimizu gave an important suggestion for conceptualization of the project, procured research equipment, reviewed the manuscript, and administered the project.

---

**Fig. S1.**
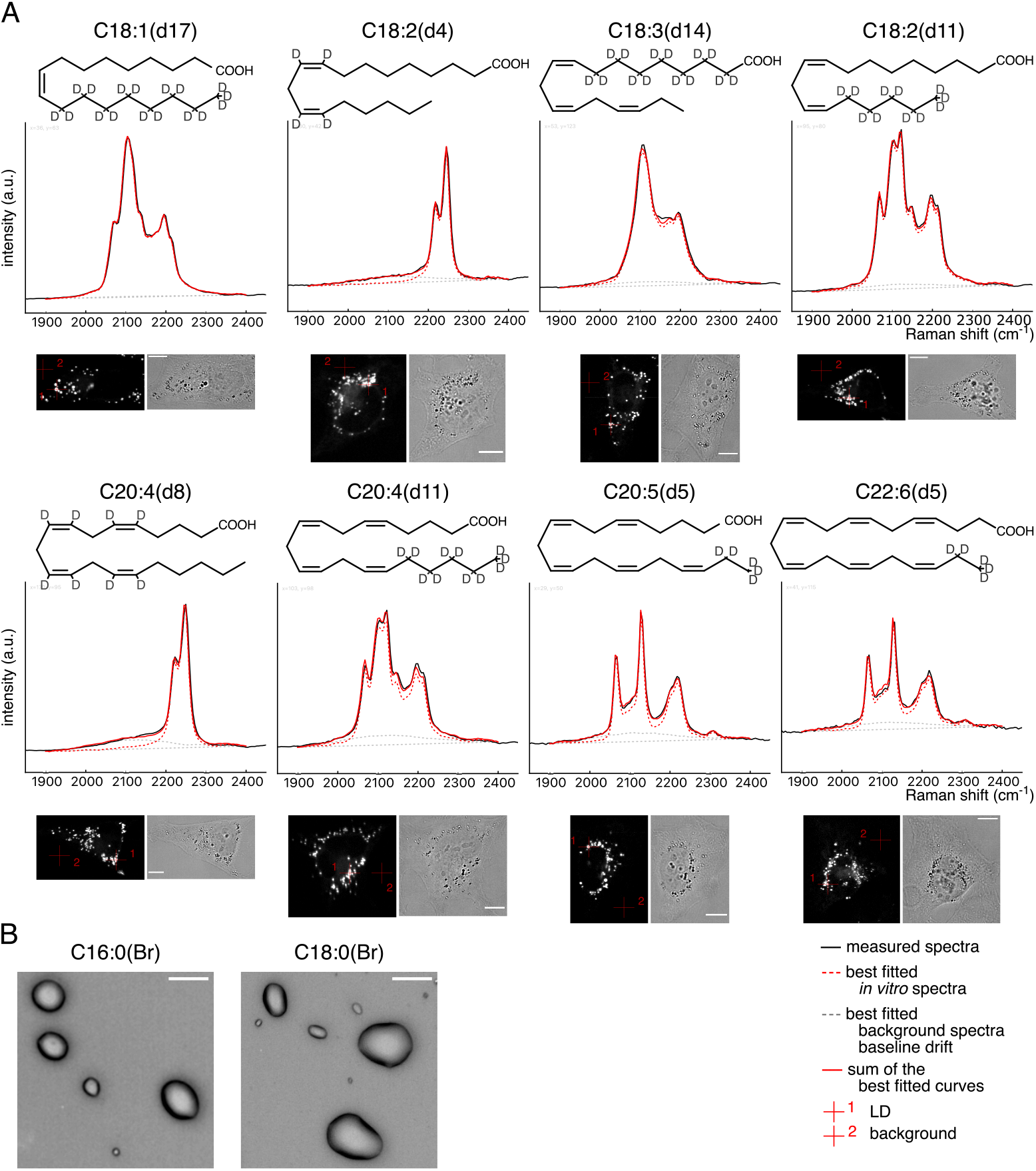
Comparison of Raman spectra of unsaturated fatty acid between *in vivo* and *in vitro*. (**A**) HeLa cells treated with indicated deuterium-labeled fatty acids (30 μM) for 24 h were fixed and observed using Raman microscopy. Representative LD spectra (represented by red crosshair 1 in the bottom images) are displayed with black lines, and the results of best fitted *in vitro* spectra, background spectra, baseline drifts, and their summations are shown in dashed red lines, dashed gray lines, dashed straight gray lines, and solid red lines, respectively. Spectra at red crosshair 2 in the bottom images were used as the background spectra for the fittings. (**B**) Bright field image of C16:0(Br) and C18:0(Br). Measurement of multiple cells and samples resulted in the similar results. Scale bars indicate 10 μm.

**Fig. S2.**
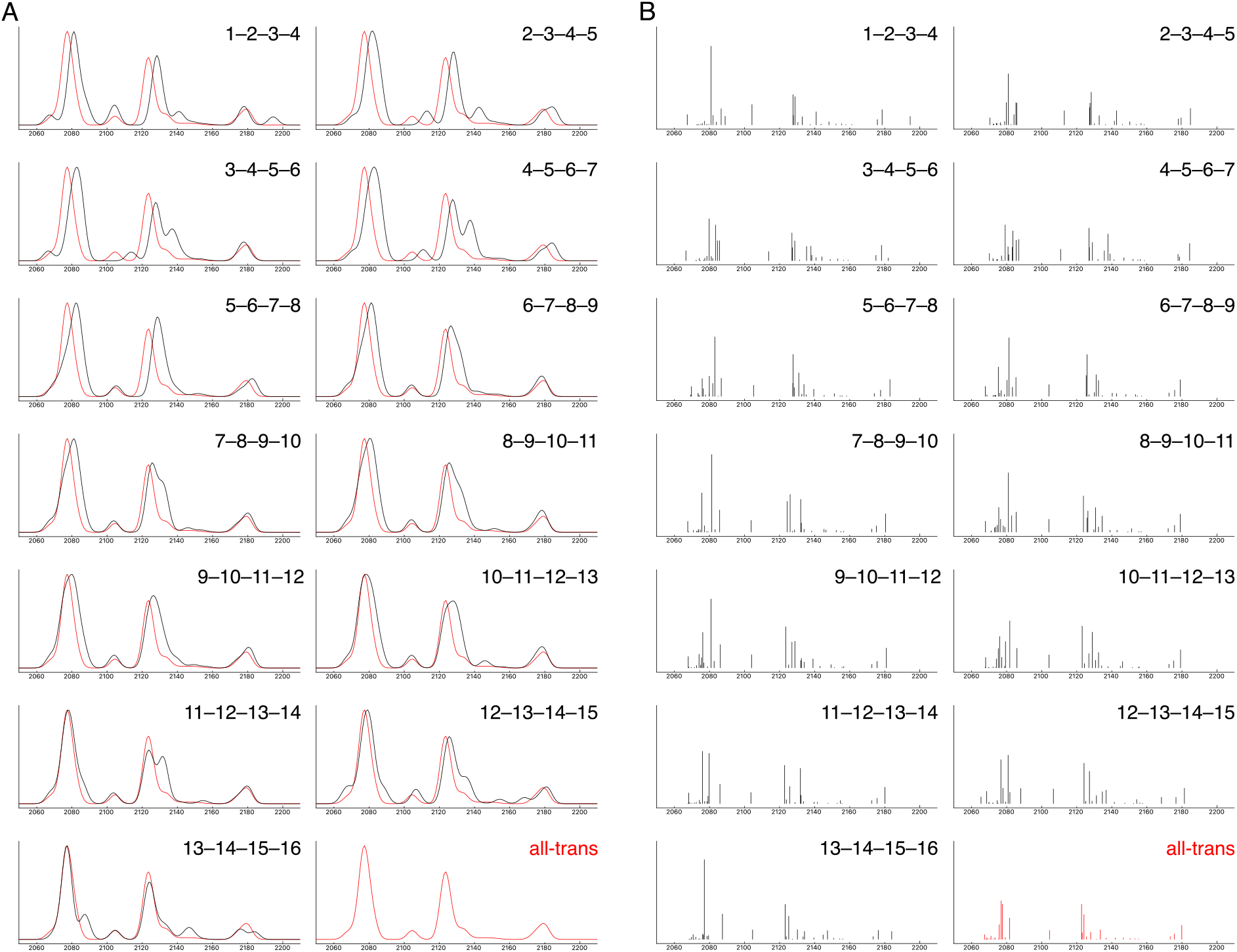
Simulated Raman spectra and Raman activity of C16:0(d31) with one gauche conformation using DFT. (**A**) The four sequential numbers at the top right of each graph indicate the positions of four sequential carbon atoms in gauche conformation. Red lines displayed in all graphs depict the simulated spectra of C16:0(d31) with all-trans conformations. Simulated Raman spectra with no gauche conformation (all-trans) and with one gauche conformation at 7-8-9-10 consecutive carbons are shown as representative in Fig. 2. (**B**) The Raman activity of corresponding Raman spectra shown in (A).

**Fig. S3.**
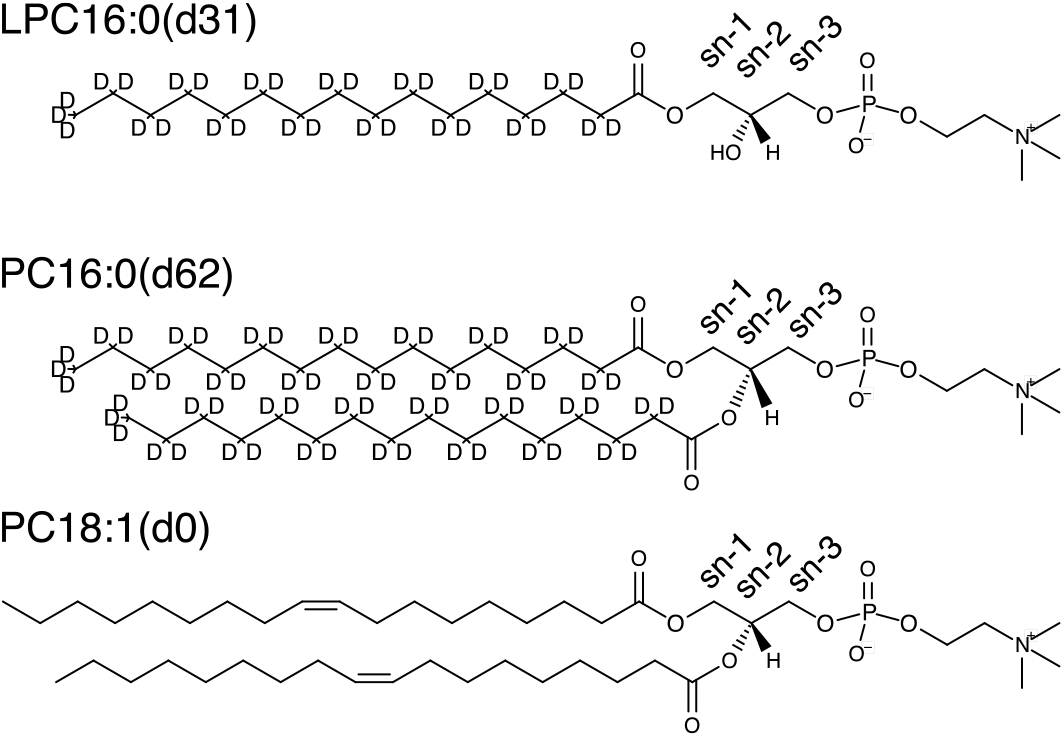
Structure of LPC16:0(d31), PC16:0(d62), and PC18:1(d0). The numbers following the letters “sn-” indicate the position of carbon atoms in the glycerol backbone.

**Fig. S4.**
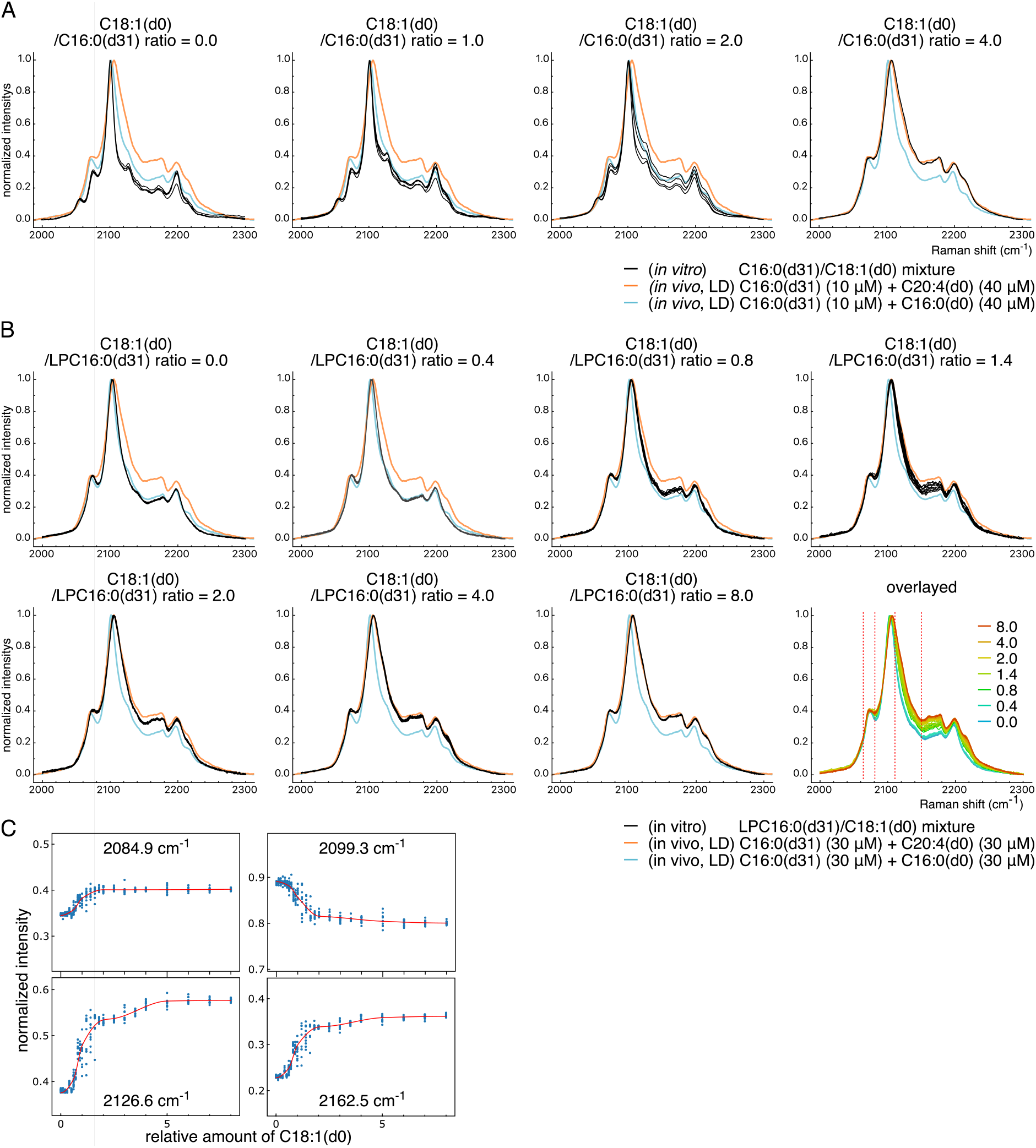
*In vitro* spectra of C16:0(d31) and LPC16:0(d31). (**A**) *In vitro* spectra of C16:0(d31) mixed with C18:1(d0). Fatty acids were mixed with the indicated ratio, and Raman spectra were then measured. For each condition, 3 to 5 spectra were measured from different locations in the sample, each displayed in the same graph with black lines. To show the similarity of *in vitro* spectra to *in vivo* spectra, *in vivo* spectra of LDs from Fig. 3A are displayed using blue and orange lines. (B) *In vitro* spectra of LPC16:0(d31) mixed with C18:1(d0). Lipids were mixed with the indicated ratio, and Raman spectra were then measured. For each condition, 9 or 10 spectra were measured from different locations in the sample, each displayed in the same graph with black lines (seven graphs except for the bottom right). To show the similarity of *in vivo* spectra to *in vitro* spectra, *in vivo* spectra of LDs from Fig. 3A are displayed using blue and orange lines. The bottom right graph depicts an overlaid graph of the other seven graphs. (**C**) Representative transitions of intensity at fixed Raman shift values. Transitions of intensity at four Raman shift values; 2084.9, 2099.3, 2126.6, and 2162.5 cm^−1^ (dashed four vertical red lines in the bottom right graphs of [B]) are displayed using blue dots. Red lines indicate interpolated curves using PCHIP algorithm.

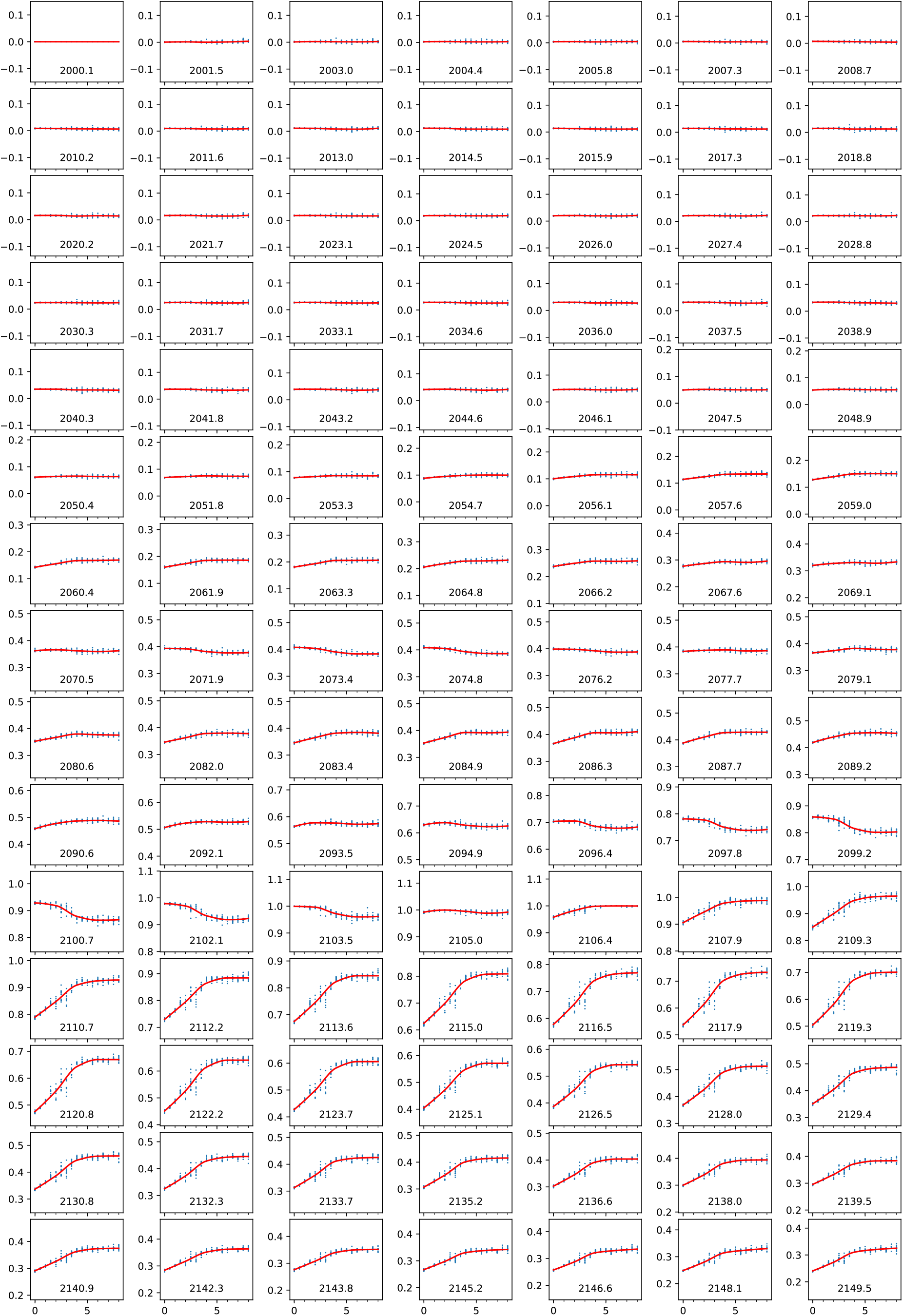

**Fig. S5.**
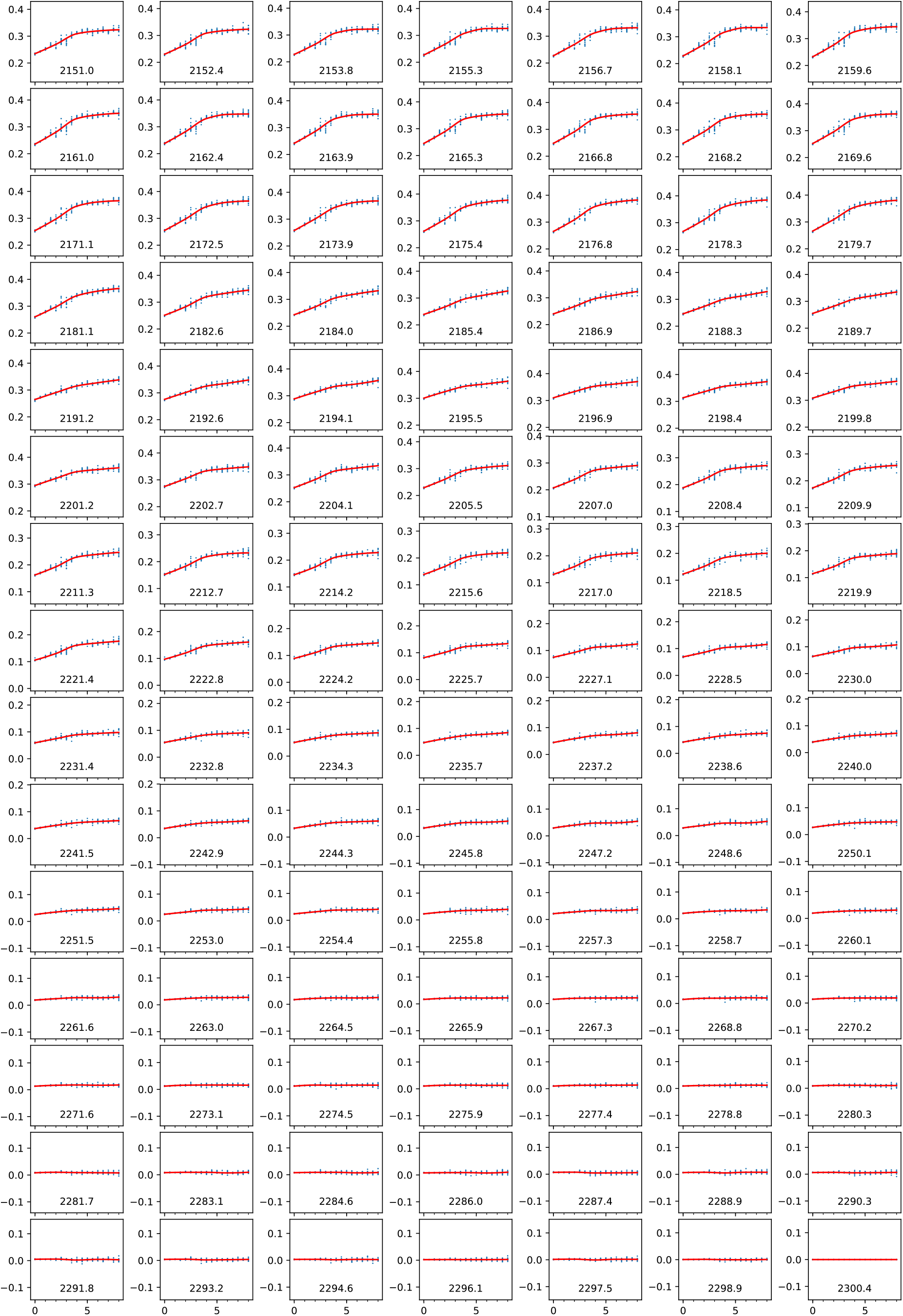
Transition of intensity at each data point with a fixed Raman shift value. Transitions of intensity at each Raman shift value (cm^−1^) are displayed using blue dots in each graph. Red lines indicate interpolated curves using PCHIP algorithm (see Materials and Methods). X- and Y-axes indicate the PC18:1(d0)/PC16:0(d31) ratio and the relative intensity at each Raman shift, respectively.

**Fig. S6.**
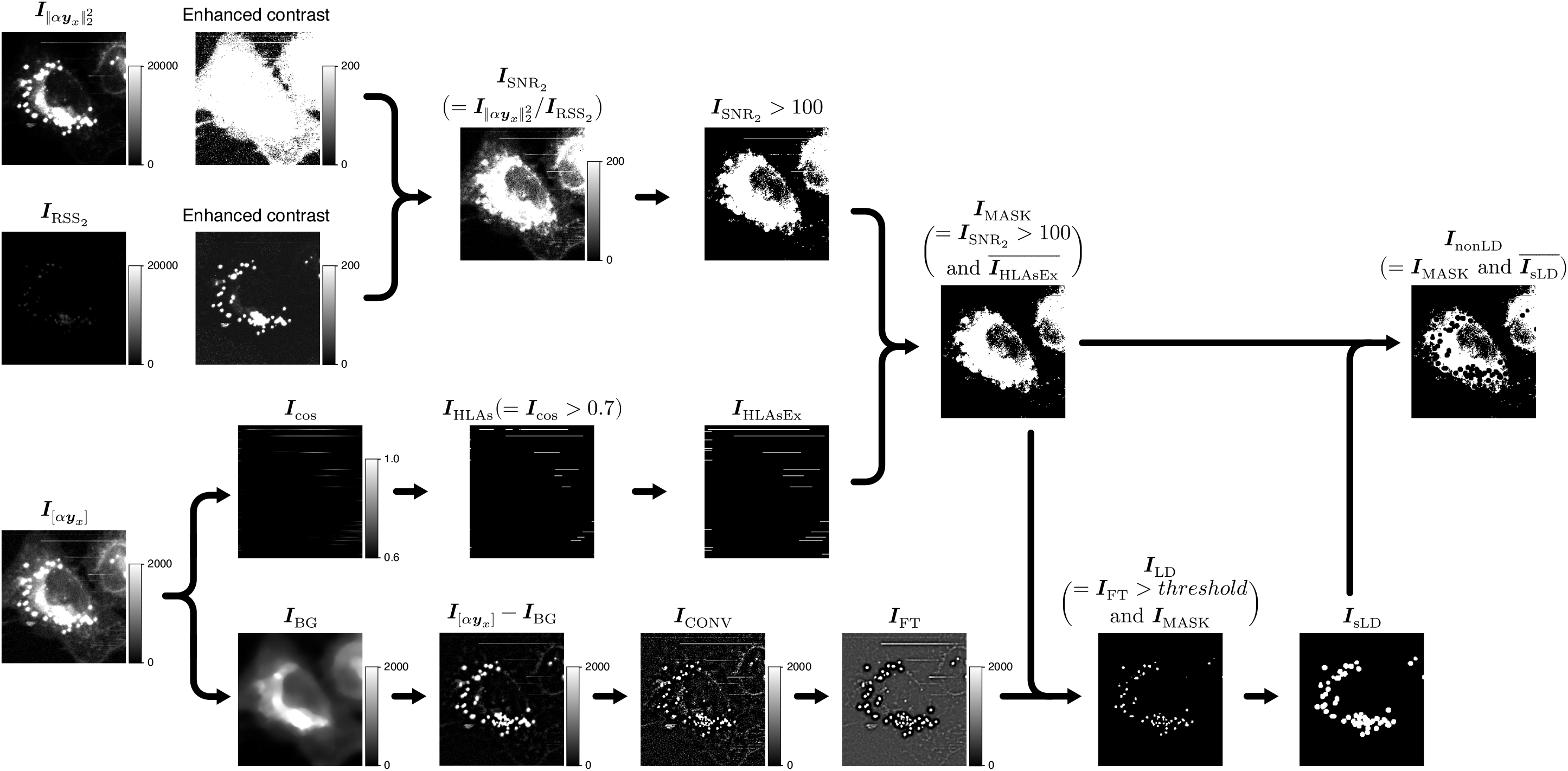
Procedure for image quantification. Image of the square sum 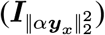, the residual sum of squares 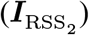, and the area under the curve 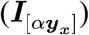 of the spectrum from C–D stretch were generated as the byproduct of converting Raman hyperspectral data into the PC18:1(d0)/PC16:0(d62) ratio image. By dividing 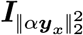 by 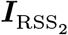, a signal to noise ratio (SNR) image was created 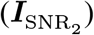, and pixels satisfying 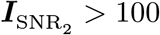 were used as a mask to omit data with low SNR. Separately, 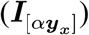 was processed for tow aims: the detection of horizontal line artifacts (HLAs), and the definition of LD regions. To detect HLAs, cosine similarity was calculated against a horizontal binary image (31 by 3), with only the middle row having an intensity of 1(***I***_cos_). Next, pixels satisfying ***I***_cos_ > 0.7 were defined as HLAs (***I***_HLAs_), followed by the horizontal extension of ***I***_HLAs_ by 10 pixels (***I***_HLAsEx_). By combining pixels with 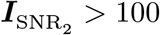 and ***I***_HLAsEx_, ***I***_MASK_ was defined and used as a mask in subsequent analyses. To define LD regions, the median background (***I***_BG_) was subtracted from 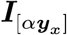, followed by convolutional processing with custom kernel ***I***_CONV_ and bandpass filter processing using Fourier transformation (***I***_FT_). Finally, a threshold defined using the modal intensity of ***I***_FT_ was applied to define LD regions (***I***_LD_). Non-LD regions were defined as the area inside ***I***_MASK_ that is not included in swelled LD regions (***I***_sLD_), which were created by swelling ***I***_LD_ by two pixels. 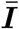 represents the inverted binary image of ***I***.

